# *GOLDEN2-like1* is sufficient but not necessary for chloroplast biogenesis in mesophyll cells of C_4_ grasses

**DOI:** 10.1101/2023.02.10.528040

**Authors:** Julia Lambret-Frotte, Georgia Smith, Jane A. Langdale

## Abstract

Chloroplasts are the site of photosynthesis. In land plants, chloroplast biogenesis is regulated by a family of transcription factors named *GOLDEN2-like* (*GLK*). In C_4_ grasses, it has been hypothesized that genome duplication events led to the sub-functionalization of *GLK* paralogs (*GLK1* and *GLK2*) to control chloroplast biogenesis in two distinct cell types: mesophyll and bundle sheath cells. Although previous characterization of *golden2* (*g2*) mutants in maize has demonstrated a role for *GLK2* paralogs in regulating chloroplast biogenesis in bundle sheath cells, the function of *GLK1* has remained elusive. Here we show that, contrary to expectations, *GLK1* is not required for chloroplast biogenesis in mesophyll cells of maize. Comparisons between maize and *Setaria viridis*, which represent two independent C_4_ origins within the Poales, further show that the role of *GLK* paralogs in controlling chloroplast biogenesis in mesophyll and bundle sheath cells differs between species. Despite these differences, complementation analysis revealed that *GLK1* and *GLK2* genes from maize are both sufficient to restore functional chloroplast development in mesophyll and bundle sheath cells of *Setaria viridis* mutants. Collectively our results suggest an evolutionary trajectory in C_4_ grasses whereby both orthologs retained the ability to induce chloroplast biogenesis but *GLK2* adopted a more prominent developmental role, particularly in relation to chloroplast activation in bundle sheath cells.

## INTRODUCTION

Chloroplasts are endosymbiotic organelles and are the site of photosynthetic reactions in a range of organisms from algae to land plants (Sagan 1967). Given the importance of chloroplast function for autotrophic growth, development of the organelle is a tightly controlled process that involves many levels of regulation (Waters and Langdale 2009). First described in the mid-1990s, the *golden2* (*g2*) mutant in *Zea mays* (maize) exhibits pale green leaves and impaired chloroplast development specifically in bundle sheath cells (Langdale and Kidner 1994, Hall *et al*. 1998). Subsequent characterization revealed that *ZmG2* encodes a transcription factor (Rossini *et al*. 2001), a founding member of the GARP transcription factor family (Riechmann *et al*. 2000). In most plant species examined, two *G2*-like (*GLK*) genes are present in the genome (Wang *et al*. 2013). *GLK* genes are defined by a characteristic helix-turn-helix DNA-binding domain related to MYB or MYB-like transcriptional factors, a conserved AREAEAA motif located immediately after the last helix and a GCT-box in the C-terminal region that is responsible for either homo- or heterodimerization (Rossini *et al*. 2001, Fitter *et al*. 2002, Safi *et al*. 2017). Functional characterization of *GLK* genes in maize and *Arabidopsis thaliana* (Arabidopsis) has shown that they are involved in the transcriptional activation of multiple photosynthesis-related genes including those encoding proteins of the light-harvesting complexes and enzymes of the chlorophyll biosynthesis pathway (Cribb *et al*. 2001, Waters *et al*. 2009). This pivotal role in chloroplast biogenesis has been associated with a myriad of physiological processes including stomatal movement, hormone signalling, defence responses and fruit ripening (Kobayashi *et al*. 2012, Nguyen *et al*. 2014, Li *et al*. 2020). The emerging picture is one where GLK proteins act as linchpins in any metabolic or developmental pathway that requires chloroplast function to be modulated.

During the evolution of land plants, the *GLK* gene family underwent multiple independent duplication events (Wang *et al*. 2013). In cases such as Arabidopsis where duplications likely occurred within species, the resulting paralogs are functionally redundant (Waters *et al*. 2009). However, duplication events in the monocots are more intriguing. The common ancestor of the Poales underwent a whole genome duplication that subdivided the *GLK* gene family into two sister clades: *GLK1* and *GLK2* (Wang *et al*. 2013). Notably, this event occurred before the many independent speciation events that led to the evolution of C_4_ photosynthesis within the Poales group (Christin and Osborne 2013). In monocot species such as rice that perform C_3_ photosynthesis, the *GLK* paralogs are functionally redundant as in Arabidopsis (Waters *et al*. 2009, Wang *et al*. 2013). Unlike C_3_ photosynthesis, where carbon reactions occur predominantly in mesophyll cell chloroplasts, in C_4_ plants carbon assimilation and fixation reactions are compartmentalized in the chloroplasts of mesophyll and bundle sheath cells, respectively (Langdale 2011). This biochemical compartmentalization is often accompanied by anatomical differences in chloroplast ultrastructure (Woo *et al*. 1970, Sage *et al*. 2011). In the C_4_ monocot species maize, *Setaria viridis* (setaria) and *Sorghum bicolor* (sorghum), *GLK1* is expressed preferentially in mesophyll cells whereas *GLK2* transcripts accumulate preferentially in bundle sheath cells, with the level of specificity differing between species (Li *et al*. 2010, Chang *et al*. 2012, Wang *et al*. 2013, John *et al*. 2014, Tausta *et al*. 2014). Notwithstanding that protein levels dictate functional capacity, defects in bundle sheath but not mesophyll chloroplast development in *g2* mutants of maize (Langdale and Kidner 1994, Cribb *et al*. 2001, Rossini *et al*. 2001) demonstrated that the preferential localization of *ZmG2* transcripts in bundle sheath cells is functionally relevant and that *GLK* paralogs in C_4_ grasses can control chloroplast development in a cell-specific manner.

*GLK* paralogs have been extensively characterized in Arabidopsis, however, little is known about gene function in monocots, and in particular the function of *GLK1* genes has not been investigated. Because *GLK1* transcripts accumulate at much higher levels in mesophyll cells than bundle sheath cells of both maize (10-fold) and setaria (30-fold) (Chang et al. 2012, John et al. 2014; Figure S1), and *Zmg2* mutants of maize show bundle sheath specific chloroplast defects, it has been hypothesized that GLK1 controls chloroplast biogenesis specifically in mesophyll cells of C_4_ grasses (Rossini *et al*. 2001). Here we test this hypothesis by generating and characterizing loss-of-function mutants in maize and setaria, two species that represent independent origins of C_4_. Our findings reveal that contrary to expectations, neither *ZmGLK1* nor *SvGLK1* are necessary for chloroplast development in mesophyll cells whereas loss of *SvGLK2* function perturbs chloroplast development in both bundle sheath and mesophyll cells.

## RESULTS

### *ZmGLK1* is not necessary for chloroplast biogenesis in mesophyll cells of maize

To determine whether *ZmGLK1* regulates chloroplast biogenesis in maize, loss-of-function mutants were generated using CRISPR/Cas9 (Figure 1A). Four independent mutant lines (referred to collectively as *Zmglk1*) were obtained, all of which exhibited the same overall growth phenotype as segregating wild-type siblings (Figure 1B, C; Figure S2). Ultrastructure analysis revealed that chloroplasts in *Zmglk1* lines were similar to wild-type, with well-developed granal thylakoids in mesophyll cells and agranal thylakoids in bundle sheath cells (Figure 1D-I; Figure S2). Total chlorophyll levels were also similar in wild-type and *Zmglk1* mutants (Figure 1J). As such, we concluded that *ZmGLK1* is not necessary for thylakoid assembly or chlorophyll biosynthesis in bundle sheath or mesophyll chloroplasts of maize.

**Figure 1.**
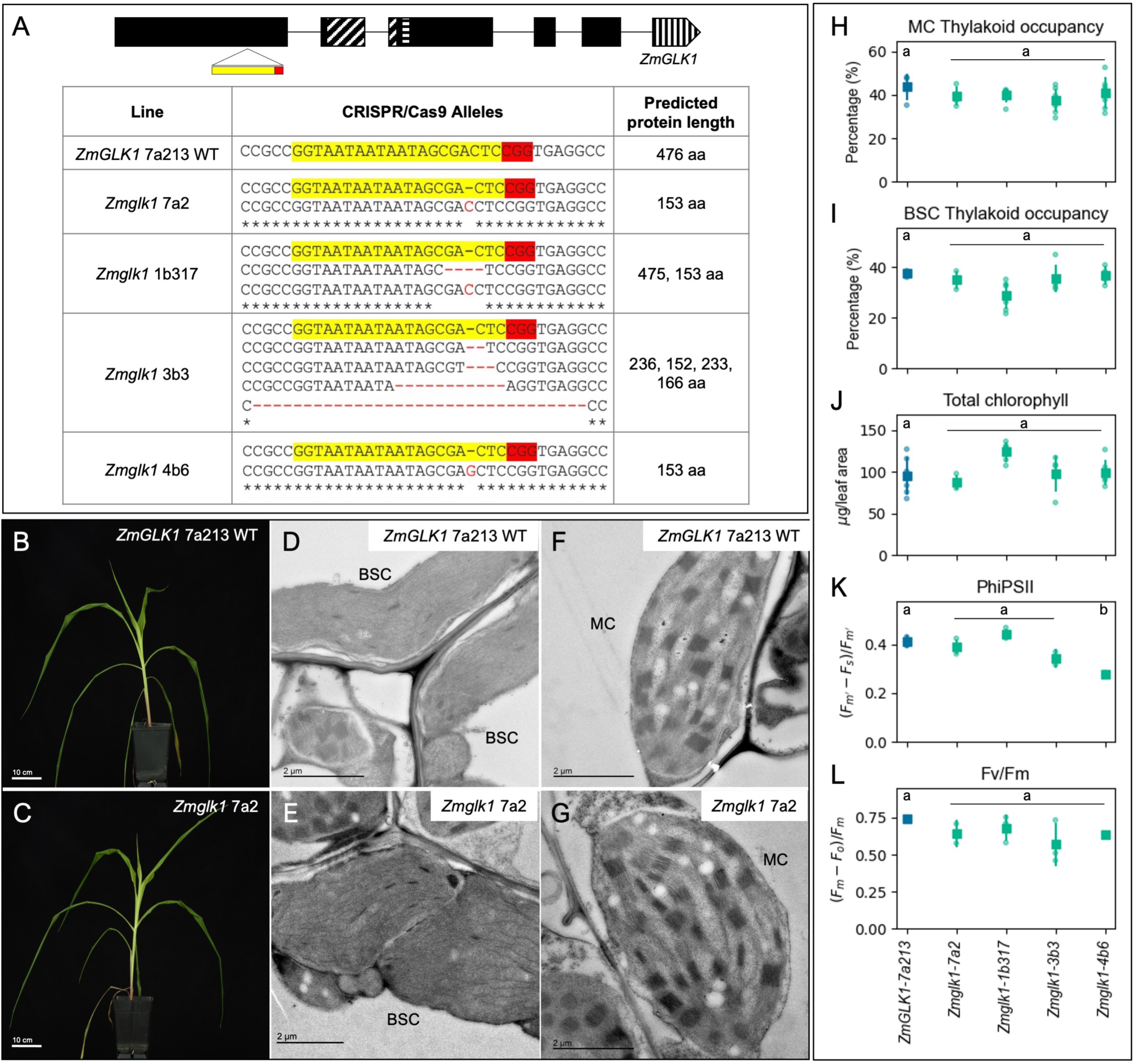
Chloroplast development is not perturbed in *Zmglk1* mutants of maize. **A)** Gene model of *ZmGLK1* showing predicted exons in black boxes and the target site of CRISPR/Cas9 mediated mutagenesis in yellow and red below. The HLH associated with MYB or MYB-related proteins is depicted by diagonal lines; the AREAEAA conserved site is highlighted with horizontal lines and the last exon that contains the GCT-box domain is depicted by vertical lines. The table shows the sequence of the wild-type (WT) and mutated alleles with the predicted protein length. The gRNA sequence is highlighted in yellow and the PAM sequence in red. **B-C)** Whole plant phenotype of wild-type (B) and a representative *Zmglk1* mutant 30 days after sowing (C). Scale bars = 10 cm. **D-G)** Electron micrographs of chloroplast ultrastructure in bundle sheath (BSC; D-E) and mesophyll (MC; F-G) cells of WT (D, F) and a representative *Zmglk1* mutant (E, G). Scale bars = 2 μm. **H-L)** Measurements of thylakoid occupancy in mesophyll cells (H), thylakoid occupancy in bundle sheath cells (I), total chlorophyll (J), PSII operating efficiency (K) and maximum quantum efficiency of PSII photochemistry (L). The WT segregant is depicted in blue and the four independent *Zmglk1* lines are depicted in green. Each dot represents a biological replicate, the square is the average value for that line and the bars are the associated standard deviations. A minimum of 5 biological replicates were evaluated for thylakoid occupancy and chlorophyll quantification, and 3 biological replicates were assessed for PSII fluorescence. Different letters indicate statistically different groups (p value ≤0.05, one-way ANOVA and Tukey’s HSD; see Table S2 for raw data).

An evaluation of chloroplast function in *Zmglk1* mutants was made by measuring photosystem II (PSII) parameters. PSII is a large protein complex comprised of the antenna and chlorophyll molecules that harvest light and together initiate the electron transfer reactions of photosynthesis (Nickelsen and Rengstl 2013). In maize, PSII accumulates in the granal thylakoids of mesophyll cells and is largely absent from the agranal bundle sheath cell chloroplasts (Woo *et al*. 1970). As such, PSII function is a proxy for mesophyll cell chloroplast function. Fluorescence measurements were therefore taken in light and dark-adapted plants to quantify PSII operating efficiency (PhiPSII) and maximum quantum efficiency of PSII photochemistry (Fv/Fm), in both wild-type and *Zmglk1* mutant plants. Figures 1K-L show that PSII is operating normally in *Zmglk1* mutants, suggesting that *ZmGLK1* is not necessary for chloroplast biogenesis or function in maize mesophyll cells.

### *SvGLK1* is not necessary for chloroplast biogenesis in mesophyll cells of setaria

Maize and setaria have different origins of C_4_ photosynthesis within the PACMAD clade (Li and Brutnell 2011) but like the orthologous *ZmGLK1*, *SvGLK1* transcripts accumulate preferentially in mesophyll cells (John *et al*. 2014). Phylogenetic analysis and amino acid sequence comparisons revealed that *SvGLK1* shares many structural features with *ZmGLK1* (Figure S3). To assess whether *SvGLK1* plays a role in chloroplast biogenesis, two independent loss-of-function mutants were generated in setaria using CRISPR/Cas9 (Figure 2A). As in maize, *Svglk1* mutants exhibited overall plant growth (Figure 2B-D) and chloroplast ultrastructure (Figure 2E-J) phenotypes that were similar to segregating wild-type siblings, with thylakoid membranes developing normally in both mesophyll and bundle sheath cells (Figure 2E-L). Furthermore, there was no significant difference in chlorophyll levels between *Svglk1* mutants and wild-type (Figure 2M), and electron flow through PSII was unperturbed (Figure 2N-O). Collectively, these results suggest that the preferential expression of *GLK1* genes in mesophyll cells of C_4_ grass species does not reflect a requirement for gene function during chloroplast development and/or maintenance.

**Figure 2.**
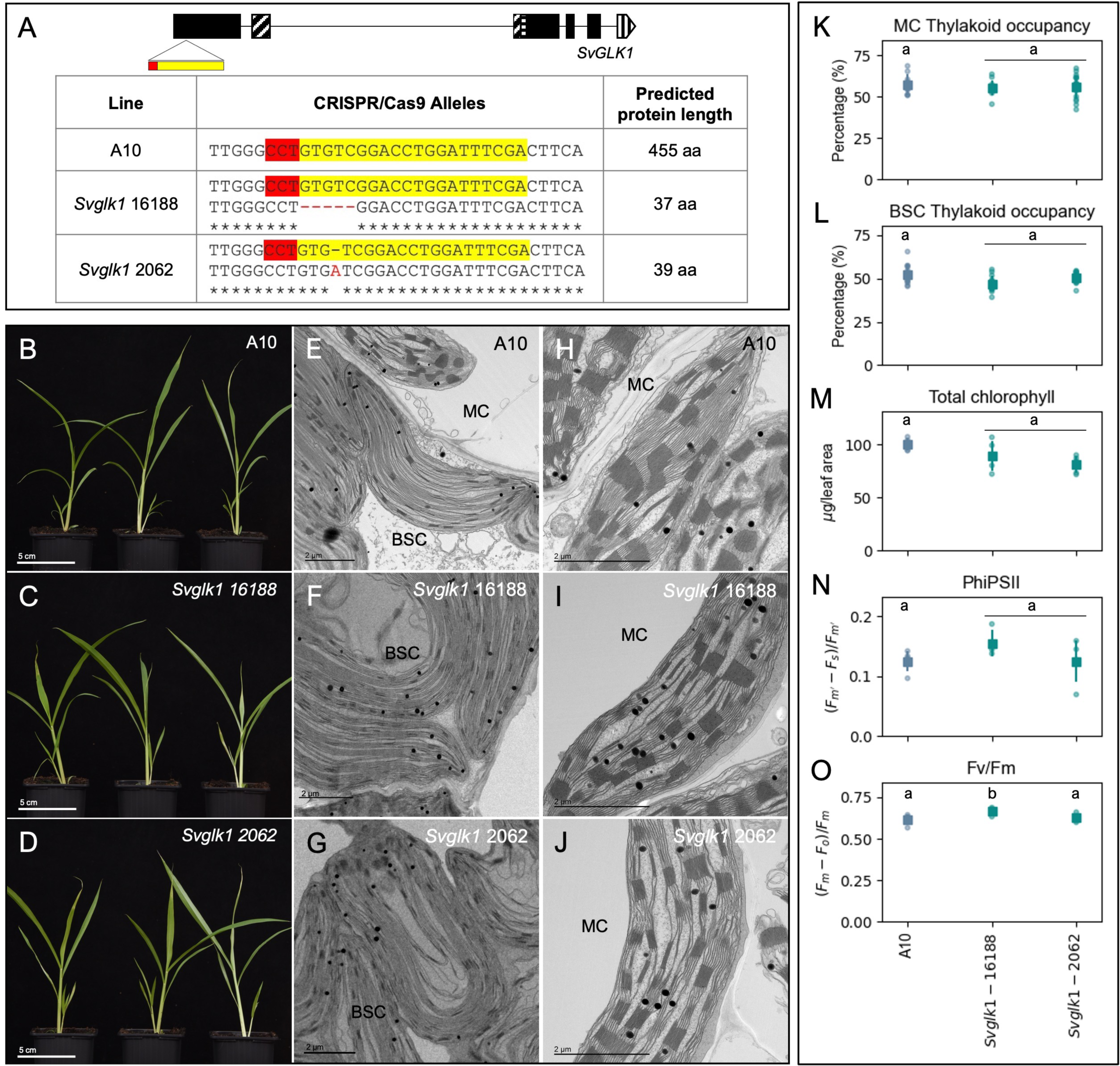
Chloroplast development is not perturbed in *Svglk1* mutants of setaria. **A)** Gene model of *SvGLK1* showing predicted exons in black boxes and the target site of CRISPR/Cas9 mediated mutagenesis in yellow and red below. The HLH associated with MYB or MYB-related proteins is depicted by diagonal lines; the AREAEAA conserved site is highlighted with horizontal lines and the last exon that contains the GCT-box domain is depicted by vertical lines. The table shows the sequence of the wild-type (WT) and mutated alleles with the predicted protein length. The gRNA sequence is highlighted in yellow and the PAM sequence in red. **B-D)** Whole plant phenotype of wild-type (B) and *Svglk1* mutants 20 days after sowing (C-D). Scale bars = 5 cm. **E-J)** Electron micrographs of chloroplast ultrastructure in bundle sheath (BSC; E-G) and mesophyll (MC; H-J) cells of WT (E, H) and *Svglk1* mutants (F, G, I, J). Scale bars = 2 μm. **K-O)** Measurements of thylakoid occupancy in mesophyll cells (K), thylakoid occupancy in bundle sheath cells (L), total chlorophyll (M), PSII operating efficiency (N) and maximum quantum efficiency of PSII photochemistry (O). The WT segregant is depicted in blue and the two independent *Svglk1* lines are depicted in green. Each dot represents a biological replicate, the square is the average value for that line and the bars are the associated standard deviations. Twelve chloroplasts were assessed for thylakoid occupancy quantification, and 5 biological replicates were assessed for chlorophyll quantification and PSII fluorescence for each sample. Different letters indicate statistically different groups (p value ≤0.05, one-way ANOVA and Tukey’s HSD; see Table S2 for raw data).

### *SvGLK2* regulates chloroplast development in both mesophyll and bundle sheath cells of setaria

Given that dimorphic chloroplast development is observed in bundle sheath and mesophyll cells of both maize and setaria, we next sought to determine whether *SvGLK2,* like its ortholog *ZmG2* (Figure S3), regulates bundle sheath chloroplast development. To this end, CRISPR/Cas9 was used to generate loss-of-function *Svglk2* mutants (Figure 3A). In contrast to maize where *g2* mutants are weaker than wild-type but are viable and fertile, any *Svglk2* mutant plants that regenerated after transformation did not progress beyond the seedling stage before dying. The only viable line recovered from tissue culture was heterozygous for *SvGLK2* and this was used to isolate a null homozygous line via self-pollination. *Svglk2* 8G5 mutants, henceforth referred to as *Svglk2*, exhibited severe growth defects, had pale yellow leaves and failed to complete the life cycle (Figure 3B). Leaf cross sections showed misshapen and smaller chloroplasts in the bundle sheath cells of *Svglk2* mutants compared to wild-type (Figure 3C-D), and observations of chloroplast ultrastructure revealed impaired chloroplast anatomy with discontinuous thylakoids that occupied less of the organelle than in wild-type (Figure 3E-F), albeit a reduction that was not statistically significant (Figure 3I). More striking, given the bundle sheath specific perturbations observed in maize *g2* mutants, was the aberrant phenotype of mesophyll chloroplasts in *Svglk2* mutants. Compared to wild-type there was a significant reduction in thylakoid membrane coverage (15% less than wild-type) and a noticeable reduction in the size and frequency of grana (Figure 3G-H, J). As in bundle sheath cells, thylakoid membranes in mutant mesophyll chloroplasts appeared fragmented (Figure 3H). The observed perturbations in bundle sheath and mesophyll cells were accompanied by a reduction in leaf chlorophyll content of more than 80% (Figure 3K). Together, the *Svglk2* and *Svglk1* mutant phenotypes suggest that SvGLK1 cannot compensate for loss of SvGLK2 function whereas SvGLK2 can compensate for loss of SvGLK1 function. As such, despite *SvGLK2* transcripts comprising only 43% of the total *GLK* transcript pool in mesophyll cells (as compared to 98% in bundle sheath cells) (John *et al*. 2014), SvGLK2 is the primary regulator of chloroplast biogenesis in both cell-types.

**Figure 3.**
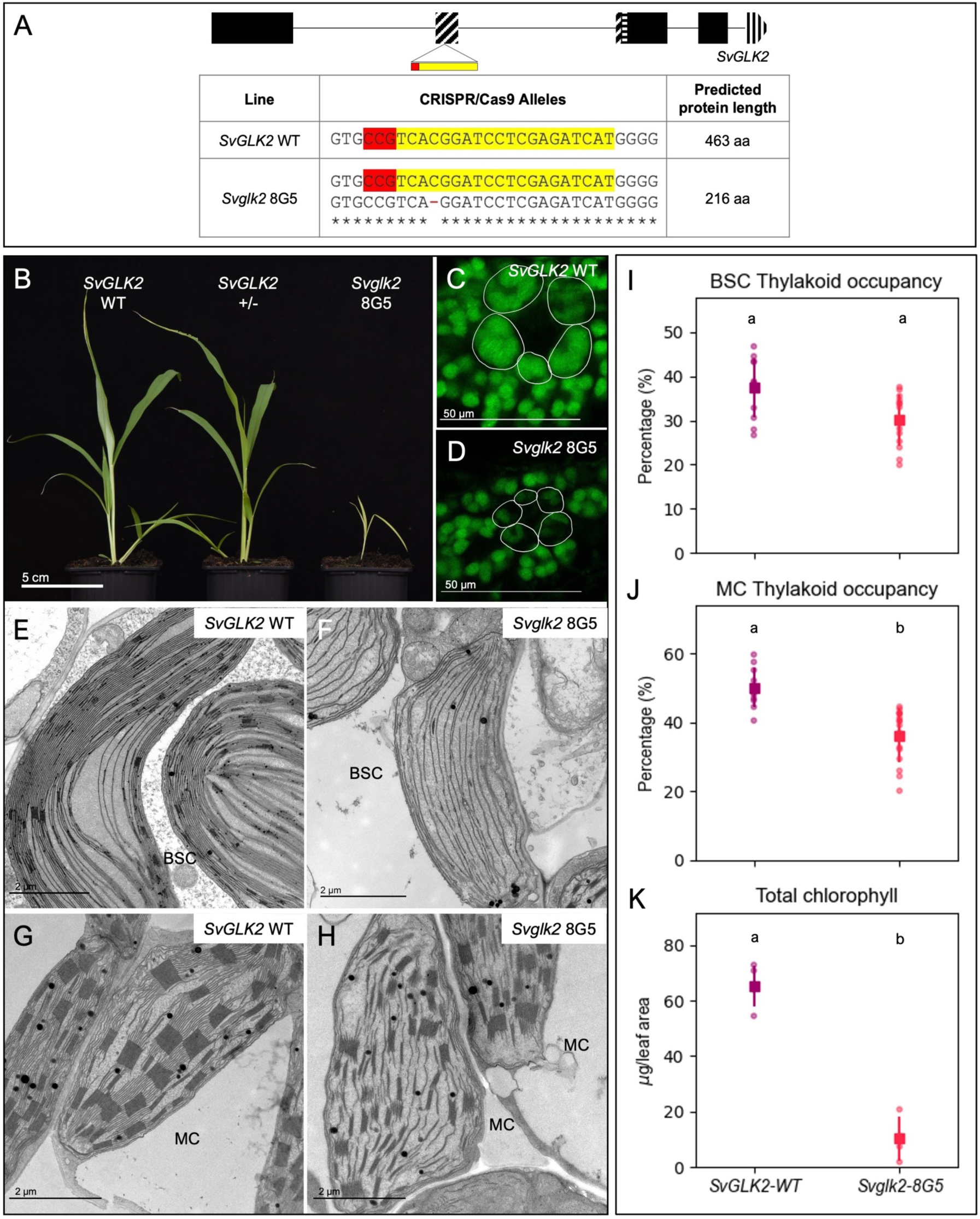
Chloroplast development is perturbed in both mesophyll and bundle sheath cells of *Svglk2* mutants of setaria. **A)** Gene model of *SvGLK2* showing predicted exons in black boxes and the target site of CRISPR/Cas9 mediated mutagenesis in yellow and red below. The HLH associated with MYB or MYB-related proteins is depicted by diagonal lines; the AREAEAA conserved site is highlighted with horizontal lines and the last exon that contains the GCT-box domain is depicted by vertical lines. The table shows the sequence of the wild-type (WT) and mutated alleles with the predicted protein length. The gRNA sequence is highlighted in yellow and the PAM sequence in red. **B)** Whole plant phenotype of wild-type, a heterozygous and a homozygous *Svglk2* mutant 20 days after sowing. Scale bar = 5 cm. **C-D)** Confocal images of leaf cross sections showing overall chloroplast structure in WT (C) and the *Svglk2* mutant (D). Bundle sheath cells are outlined in white. Scale bars = 50 μm. **E-H)** Electron micrographs of chloroplast ultrastructure in bundle sheath (BSC; E-F) and mesophyll (MC; G-H) cells of WT (E, G) and the *Svglk2* mutant (F, H). Yellow arrows point to fragmented membranes. Scale bars = 2 μm. **I-K)** Measurements of thylakoid occupancy in bundle sheath cells (I), thylakoid occupancy in mesophyll cells (J) and total chlorophyll (K). The WT segregant is depicted in purple and the *Svglk2* line is depicted in pink. Each dot represents a biological replicate, the square is the average value for that line and the bars are the associated standard deviations. Twelve chloroplasts were assessed for thylakoid occupancy quantification, and 5 biological replicates were assessed for chlorophyll quantification and PSII fluorescence for each sample. Different letters indicate statistically different groups (p value ≤0.05, one-way ANOVA and Tukey’s HSD; see Table S2 for raw data).

### *GLK2* transcript levels are not upregulated in *glk1* mutants of maize or setaria

Given that *ZmGLK1* and *SvGLK1* transcripts represent 95% and 57% of the total *GLK* transcript pool in mesophyll cells of the respective species (Chang et al. 2012, John et al. 2014; Figure S1), the absence of chloroplast defects in this cell-type in loss of function mutants is intriguing. It is particularly notable because all *GLK* genes characterized to date have a role in chloroplast biogenesis, albeit a redundant role alongside paralogs in many cases (Fitter *et al*. 2002, Wang *et al*. 2013). Even a redundant role is unlikely in setaria given the *Svglk2* mutant phenotype. However, redundancy could explain the lack of a *Zmglk1* mutant phenotype if *ZmG2* transcripts are translated very efficiently in mesophyll cells of maize. Alternatively, loss of *ZmGLK1* function could lead to altered regulation of *ZmG2* expression and/or to the upregulation of genes encoding the GATA transcription factors GNC and CGA1 which also regulate chloroplast development (Chiang *et al*. 2012, Hudson *et al*. 2013) and have been shown to partially rescue loss of GLK function in Arabidopsis (Zubo *et al*. 2018). To test this hypothesis, transcript levels of *ZmGLK1*, *ZmG2*, *ZmGNC* and *ZmCGA1* were quantified in maize using the *Zmglk1* gene edited mutants generated above alongside the previously reported transposon-induced *Zmg2-bsd1-s1* mutant (Cribb *et al*. 2001). In each case the mutant allele produces transcripts that encode non-functional protein(s). Figure 4 reveals that loss of *Zmglk1* function does not lead to consistently elevated levels of *ZmGLK1* (Figure 4A), *ZmG2* (Figure 4B), *ZmGNC* (Figure 4C) or *ZmCGA1* (Figure 4D) transcripts. A small but significant increase in *ZmG2*, *ZmGNC* and *ZmCGA1* transcript levels was seen in one of the four *Zmglk1* mutant lines but that cannot explain how chloroplasts develop normally in all four lines. We thus conclude that transcription of *ZmG2, ZmGNC* and *ZmCGA1* is not upregulated in mesophyll cells when *Zmglk1* function is lost.

**Figure 4.**
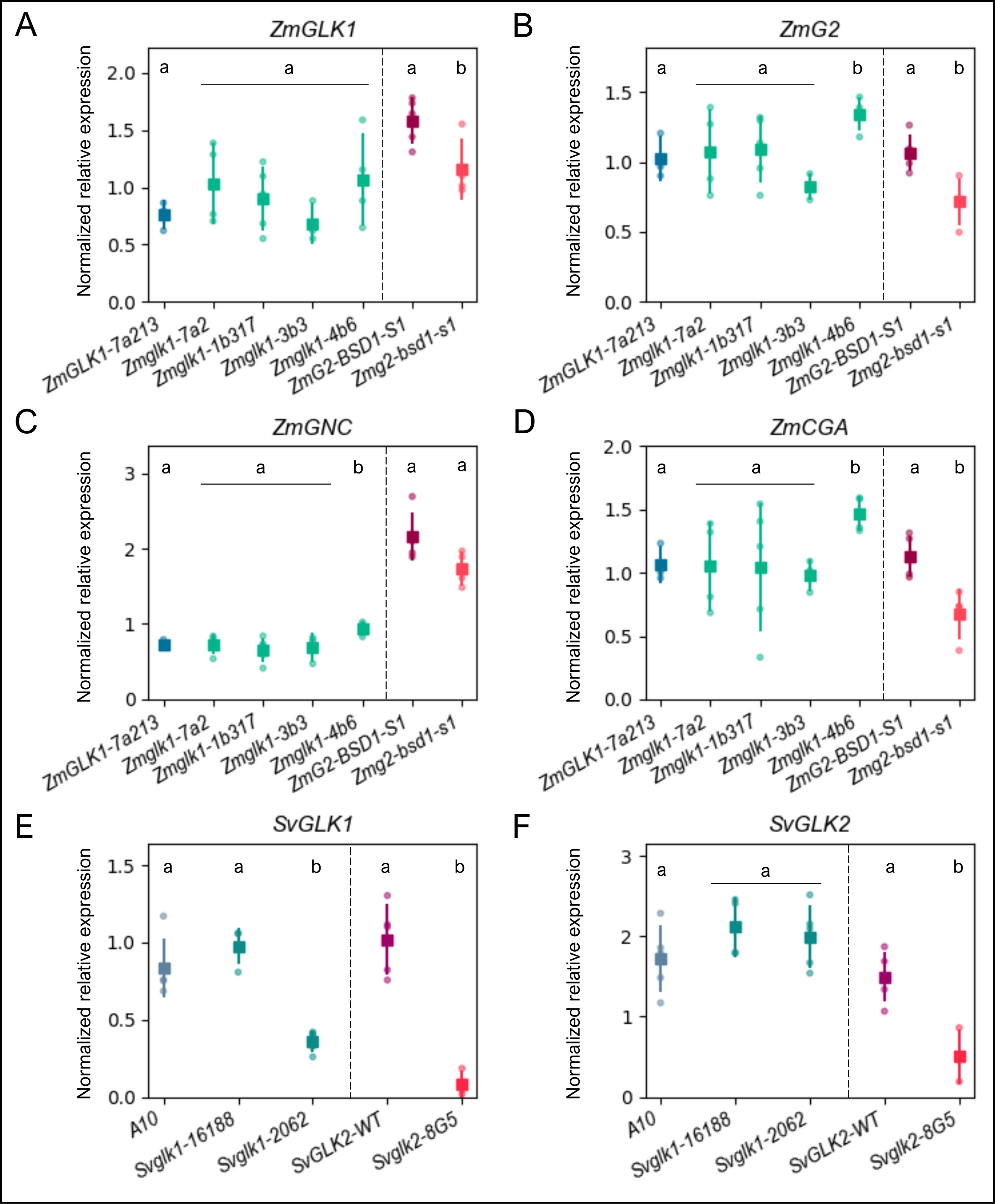
*GLK1* and *GLK2* transcript levels are reduced in *Svglk2* and *Zmg2* mutants of setaria and maize. **A-H)** Quantitative analysis of gene expression via qPCR in segregating wild-type (WT) siblings and *glk* mutants of maize (A-D) and setaria (E-F). Relative normalized expression of *ZmGLK1* (A), *ZmG2* (B), *ZmGNC* (C), *ZmCGA1* (D), *SvGLK1* (E) and *SvGLK2* (F) was calculated based on the expression of two reference genes for each sample. Each dot represents a biological replicate, the square is the mean value and the bars are the associated standard deviations. A minimum of 3 and 5 biological replicates were evaluated for maize and setaria samples, respectively. Statistical significance was calculated using a Student’s t-test and pairwise comparison with the respective WT sample. For each species comparison, different letters indicate statistically different groups (p value ≤0.05; see Table S2 for raw data).

Whereas loss of ZmGLK1 function in maize does not impact on *ZmGLK1* or *ZmG2* transcript levels, loss of ZmG2 function in bundle sheath cells leads to a statistically significant reduction in the accumulation of both *ZmGLK1* and *ZmG2* transcripts (Figure 4A, B), and a similar situation is seen in setaria where both *SvGLK1* and *SvGLK2* transcript levels are dramatically reduced in *Svglk2* mutants (Figure 4E, F). This observation suggests that the ZmG2/SvGLK2 orthologs, either directly or indirectly, promote the accumulation of both *GLK1* and *GLK2* transcripts in the respective species. Consistent with this suggestion, multiple G-box elements which are bound by Arabidopsis GLK proteins (Waters *et al*. 2009) are present in *ZmGLK1*, *ZmG2* and *SvGLK2* promoter sequences (Figure S4), and a GLK binding site (Tu *et al*. 2022) is present 1840 bp upstream of the ATG in the *SvGLK1* gene. Reduced levels of *SvGLK1* transcripts in one of the two *Svglk1* mutant lines (Figure 4E) suggest that *SvGLK1* might regulate its own transcript levels via this binding site but the absence of such an effect in the other mutant line makes it equally likely that the specific edits in line #2062 lead to transcript turnover, possibly via the nonsense-mediated decay pathway. Collectively these data suggest that chloroplast development in C_4_ grasses is predominantly regulated by genes of the *GLK2* clade.

### *ZmGLK1* and *ZmG2* are both sufficient to restore dimorphic chloroplast development in mesophyll and bundle sheath cells of *Svglk2* mutants

Our findings thus far show that *SvGLK2* and *ZmG2* play a more central role in controlling chloroplast biogenesis in C_4_ grasses than their respective *GLK1* paralogs. To evaluate whether this difference in necessity also applies to sufficiency, *ZmGLK1* and *ZmG2* were expressed under the control of the constitutive maize ubiquitin gene promoter (*ZmUBI*_pro_) in the chloroplast-defective *Svglk2* mutant background. Three independent lines were obtained for each construct and quantitative analysis of transgene transcript levels revealed differences between lines that correlated directly with plant phenotype (Figure 5A-D). For example, the weaker plants in line #17808-15254 accumulated around half the level of *ZmGLK1* transcripts seen in healthier plants of the other two lines (Figure 5A, 5C). Notably plants of all six lines were viable, healthy and fertile, suggesting complementation of the *Svglk2* mutant phenotype (compare *Svglk2* plants in Figure 3B with Figure 5A, B).

**Figure 5.**
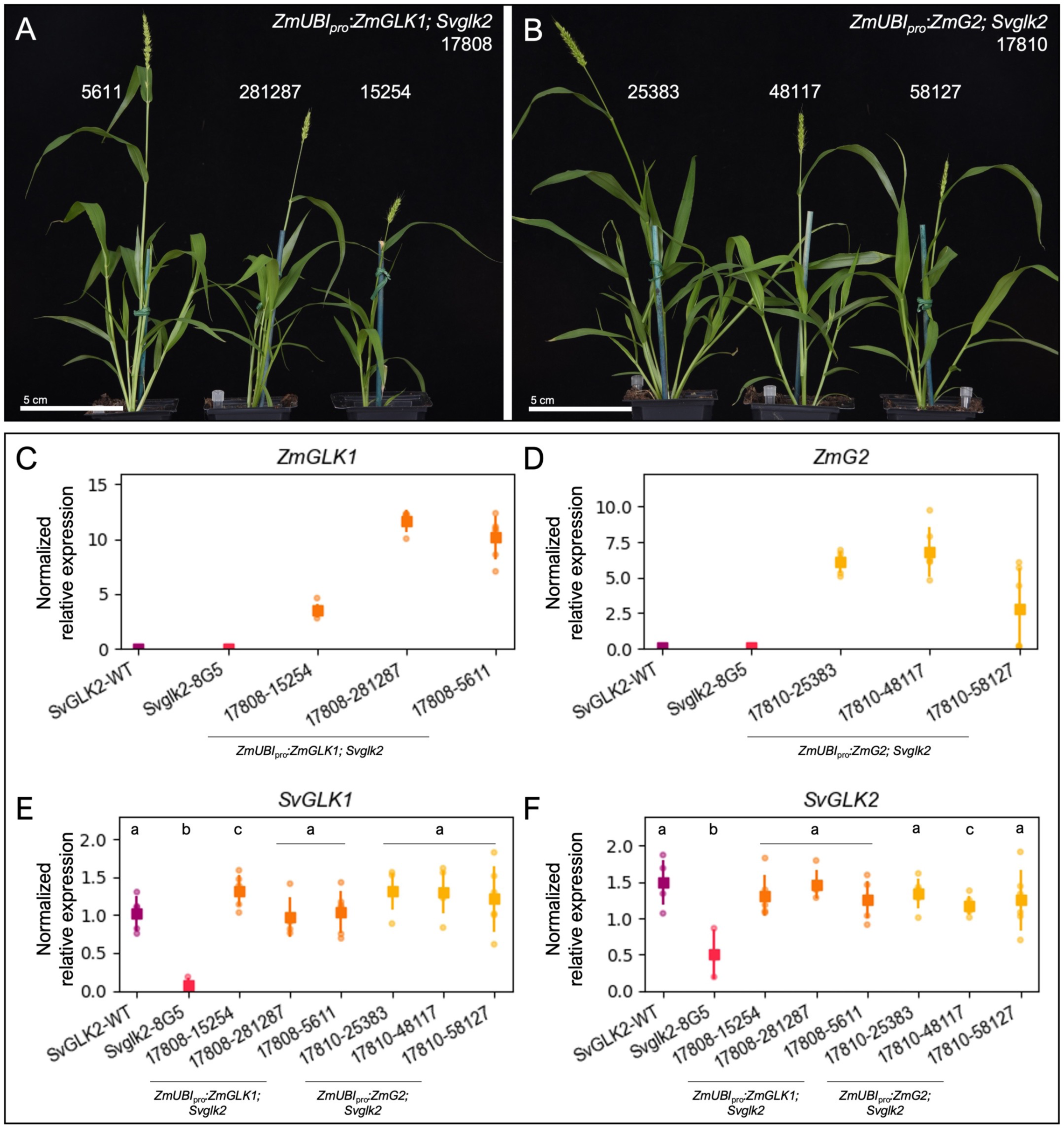
*ZmGLK1* and *ZmG2* complement the *Svglk2* phenotype and activate transcription of the setaria homologs. **A-B)** Whole plant phenotype of *Svglk2* mutant overexpressing *ZmGLK1* (A) or *ZmG2* (B) 30 days after sowing. Scale bars = 5 cm. **C-F)** Quantitative analysis of gene expression via qPCR of *ZmGLK1* (C), *ZmG2* (D), *SvGLK1* (E) and *SvGLK2* (F). Relative gene expression was calculated based on the expression of two reference genes for each sample. Each dot represents one of the 5 biological replicates evaluated, the square is the mean value and the bars are the associated standard deviations. Statistical significance was calculated using a Student’s t-test and pairwise comparison with the respective wild-type (WT) sample. Different letters indicate statistically different groups (p value ≤0.05; see Table S2 for raw data).

Given that loss of *Svglk2* function leads to reduced levels of both *SvGLK1* and *SvGLK2* transcripts (Figure 4E, F), we next sought to evaluate whether ZmGLK1 or ZmG2 could influence endogenous *SvGLK1* or *SvGLK2* expression. Figure 5E, F show that *SvGLK1* and *SvGLK2* transcripts were restored to wild-type levels in *Svglk2* lines expressing either *ZmGLK1* or *ZmG2* transgenes. In fact, regardless of the expression level of either *ZmGLK1* or *ZmG2*, all complemented lines exhibit the same wild-type-like level of *SvGLK1* and *SvGLK2* expression (Figure 5C-F). Both ZmGLK1 and ZmG2 can thus activate expression of *SvGLK1* and *SvGLK2*, providing further support for the existence of a regulatory network between *GLK* genes and for the likely relevance of G-box and GLK binding motifs in the promoter regions (Figure S4). The functional significance of this observation must be considered in the context of transgene expression driven by a constitutive promoter and gene-edited *Svglk2* transcripts encoding non-functional proteins (and hence any effect on *SvGLK1* expression is a direct effect of the maize transgene and not via *SvGLK2*). In this regard, we must deduce that the transgenic lines expressing *ZmGLK1* accumulate transcripts encoding functional ZmGLK1 and SvGLK1 proteins in both mesophyll and bundle sheath cells, and those expressing *ZmG2* accumulate transcripts encoding functional ZmG2 and SvGLK1 proteins, similarly in both cell types.

To determine whether *ZmGLK1* and/or *ZmG2* can rescue chloroplast ultrastructure defects in mesophyll and/or bundle sheath cells of *Svglk2* mutants, confocal images and transmission electron micrographs of transverse leaf sections from all six transgenic lines were examined. In all cases, chloroplasts were restored to normal size (Figure 6A-C, Figure S5), with predominantly agranal thylakoids in bundle sheath cell chloroplasts (Figure 6D-F, Figure S5) and granal thylakoids in mesophyll chloroplasts (Figure 6G-I, Figure S5). Quantification revealed that with the exception of the one line that had low levels of transgene expression (#17808-15254), thylakoid occupancy levels in mesophyll (Figure 6J) and bundle sheath (Figure 6K) chloroplasts of transgenic lines were statistically equivalent (or in one case higher) than in wild-type plants. Consistent with ectopic expression of *GLK* genes, mature chloroplasts were also observed in the leaf vasculature of all complemented lines, more with *ZmG2* than *ZmGLK1* (Figure 6A-C, Figure S5). Irrespective of which maize ortholog was overexpressed, the total chlorophyll content in leaves of transgenic plants was restored to wild-type levels (Figure 6L). Notwithstanding the fact that SvGLK1 may be functional in bundle sheath and mesophyll cells of all transgenic plants, collectively these data demonstrate that ZmGLK1 and ZmG2 proteins are functionally equivalent in the context of the C_4_ setaria leaf and that the dimorphic identity of bundle sheath and mesophyll cell chloroplasts develops regardless of which maize ortholog is overexpressed.

**Figure 6.**
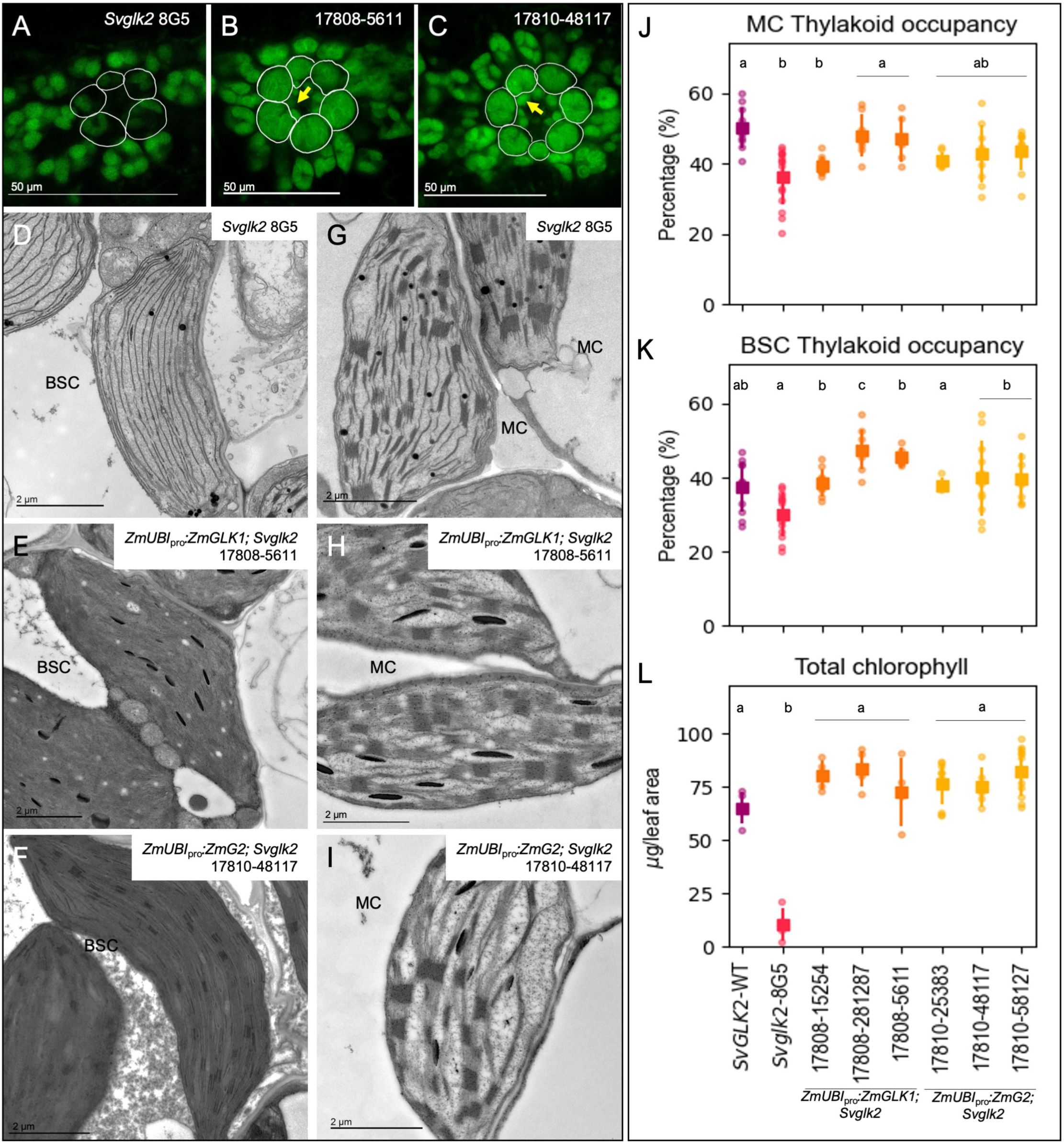
*ZmGLK1* and *ZmG2* are sufficient to induce dimorphic chloroplast development in both mesophyll and bundle sheath cells of setaria. **A-C)** Confocal images of leaf cross sections showing overall chloroplast morphology in the *Svglk2* mutant (A, same as Figure 3D), and representative lines expressing *ZmGLK1* (B) or *ZmG2* (C). Bundle sheath cells are outlined in white. Yellow arrows indicate ectopic chloroplasts developing in vascular cells. Scale bars = 50 μm. **D-I)** Electron micrographs of chloroplast ultrastructure in bundle sheath (BSC; D-F) and mesophyll (MC; G-I) cells of the *Svglk2* mutant (D-G; same as Figure 3G, H) and one representative line overexpressing *ZmGLK1* (E-H) or ZmG2 (F-I). Scale bars = 2 μm. **J-L)** Measurements of thylakoid occupancy in mesophyll cells (J), thylakoid occupancy in bundle sheath cells (K) and total chlorophyll (L). The wild-type (WT) segregant is depicted in purple, the *Svglk2* line in pink, the three independent lines overexpressing *ZmGLK1* in orange and those overexpressing *ZmG2* in yellow. Each dot represents a biological replicate, the square is the average value for that line and the bars are the associated standard deviations. A minimum of 10 chloroplasts were evaluated for thylakoid occupancy quantification and 5-10 biological replicates were used for chlorophyll quantification. Different letters indicate statistically different groups in (p value ≤0.05, one-way ANOVA and Tukey’s HSD; see Table S2 for raw data).

## DISCUSSION

Prior to this study it was known that *GLK* genes are duplicated in many land plant species and that in species that carry out C_3_ photosynthesis the duplicate genes act redundantly to regulate chloroplast biogenesis in all photosynthetic cells (Waters *et al*. 2008, Bravo-Garcia *et al*. 2009, Wang *et al*. 2013). It had also been hypothesized that *GLK1* and *GLK2* genes regulate dimorphic chloroplast development in the two photosynthetic cell-types of C_4_ species, with each gene functioning in one of the two cell-types (Rossini *et al*. 2001). This hypothesis was based on the observation that *GLK1* genes are preferentially expressed in mesophyll cells of maize, sorghum and setaria whereas *GLK2/G2* genes are preferentially expressed in bundle sheath cells (Li *et al*. 2010, Chang *et al*. 2012, John *et al*. 2014, Wang *et al*. 2014), and loss of *Zmg2* function in maize perturbs bundle sheath but not mesophyll chloroplast development (Cribb *et al*. 2001). Through the phenotypic characterization of *glk1* loss-of-function mutants in maize and setaria (Figures 1, 2 & 4), *glk2* loss-of-function mutants in setaria (Figures 3 & 4), and *glk2* setaria mutants complemented with maize *GLK1* and *GLK2* genes (Figure 5 & 6), we have revealed a more complex situation both with respect to the developmental processes operating within individual C_4_ species and to likely evolutionary trajectories.

### GLK proteins have equivalent functional capacity

*GLK* genes duplicated in a common ancestor of the Poales, before the speciation of maize, sorghum, setaria and rice, and before the evolution of C_4_ photosynthesis (Wang et al. 2013). Given that *GLK* genes regulate chloroplast biogenesis in the moss *Physcomitrium patens* (Yasumura *et al*. 2005, Bravo-Garcia *et al*. 2009), which last shared a common ancestor with flowering plants over 450 million years ago (Kenrick and Crane 1997, Wickett *et al*. 2014), it is reasonable to assume that prior to this duplication the ancestral *GLK* gene played a role in chloroplast development. The duplication event, which must have occurred between 85-95 million years ago (Gallaher *et al*. 2022), provided an opportunity for neo- and/or sub-functionalization of gene activity and thus for divergent roles to evolve in different grass species. It has previously been suggested that the compartmentalized expression of *GLK1* and *GLK2* genes in mesophyll and bundle sheath cells of C_4_ species reflects distinct roles in the biogenesis of granal versus agranal chloroplasts. This hypothesis was supported by the observation that *GLK* homologs are not differentially expressed in *Gynandropsis gynandra* (formerly known as *Cleome gynandra*), a C_4_ dicot that exhibits granal thylakoids in chloroplasts of both mesophyll and bundle sheath cells (Marshall *et al*. 2007, Wang *et al*. 2013). However, data presented here show that appropriate cell-type specific thylakoid stacking is restored in mesophyll and bundle sheath cells of *Svglk2* mutants when either *ZmGLK1* or *ZmG2* is constitutively expressed (Figures 5 & 6). These results demonstrate that the two maize proteins have equivalent functional capacity. Furthermore, the data are consistent with a recent report which showed that the identity of genes activated by GLK transcription factors is determined by *cis* regulatory factors, such that a heterologous GLK protein will activate downstream targets that are normally activated by endogenous GLK proteins in the host species as opposed to targets that are activated by the heterologous protein in its species of origin (Tu *et al*. 2022). Together these observations suggest that the evolution of dimorphic chloroplasts in C_4_ species was unlikely to have been mediated by sub-functionalization of GLK protein activity.

### Despite equivalent functional capacity, GLK2 plays a dominant role in both maize and setaria

The difference in the proportion of *GLK2:GLK1* transcripts in mesophyll cells of maize (5:95) versus setaria (43:57) can partially explain the different phenotypes observed in loss-of-function mutants. In maize, where relative levels of *ZmG2* are only 5% in mesophyll cells, *Zmg2* mutants exhibit perturbed chloroplast development only in bundle sheath cells (where *ZmG2* transcripts comprise 65% of the total GLK pool). By contrast, relative *GLK2* levels of 43% in mesophyll and 98% in bundle sheath cells of setaria are associated with defective chloroplast development in both cell-types (Figure 3). This observation was initially counter-intuitive because it would be expected that, as in C_3_ plants, the high levels of *SvGLK1* transcripts in mesophyll cells would compensate for loss of SvGLK2 function. However, we observed that *SvGLK1* transcript accumulation was virtually abolished in *Svglk2* mutants and levels of the non-functional *Svglk2* transcripts were also reduced (Figure 4). As such, we conclude that SvGLK2 activates expression (or transcript stabilization) both of itself and of *SvGLK1,* and that the *Svglk2* mutant phenotype reflects loss of function of both genes. Collectively, these data suggest that SvGLK2 is the primary regulator of chloroplast development in both bundle sheath and mesophyll cells of setaria.

Whereas normal mesophyll chloroplast development in the absence of GLK1 function can be explained in setaria because SvGLK2 likely compensates for loss of GLK1 function, the mutant phenotype in maize was unexpected. In every land plant species examined to date, a role for *GLK* genes in chloroplast development has been revealed, albeit often a redundant role with paralogs operating in the same cell-type. Because ZmGLK1 is highly expressed in maize mesophyll cells and no other GLK-like transcripts appear as possible candidates for a redundant paralog, a previously unknown mechanism must be operating in this cell-type. Although GLK gene function has previously been shown to be cell-autonomous in Arabidopsis, it is possible that a non-cell-autonomous mechanism could operate in maize. Indeed, there is some evidence of this because *ZmGLK1* transcript levels are reduced when G2 function is lost, even though *G2* transcript levels in wild-type mesophyll cells are only 5% of the GLK pool (Figure 4). In such a scenario, loss of GLK1 function in mesophyll cells would be compensated for by direct or indirect activity of G2 proteins in the bundle sheath cells. Any such effect would only be revealed in double *Zmglk1;g2* mutants, which would have defective chloroplasts in both cell-types. Regardless of whether ZmG2 compensates for loss of ZmGLK1 function, the GLK2 paralog plays the dominant role in chloroplast biogenesis in maize, as in setaria.

### *cis*-regulatory sequences are more variable in *GLK2* than *GLK1* genes

Given that *GLK* gene paralogs encode functionally equivalent proteins both in C_3_ species where they act redundantly and in the C_4_ species maize, the question arises of why the paralogs are expressed in a cell-type preferential manner in C_4_ grass species (at least in the ones that have been examined). Estimates based on the topology of BEP (C_3_ species only) and PACMAD (C_3_ & C_4_ species) clades in the Poales suggest that maize diverged from rice ∼80 million years ago (Gallaher *et al*. 2022) and that within the PACMAD clade, maize and sorghum share a common history of C_4_ evolution whereas independent events occurred in maize and setaria around ∼45 million years apart (Gallaher *et al*. 2022). Both *GLK* gene paralogs are predominantly expressed in mesophyll cells of C_3_ rice leaves, with *GLK1* transcripts accumulating at higher levels than *GLK2* (Hua *et al*. 2021), and gene function being redundant (Wang *et al*. 2013). Notably, both transcripts can also be detected in bundle sheath cells, at levels roughly equivalent to those of *GLK2* in mesophyll cells, suggesting that the few chloroplasts that develop in rice bundle sheath cells are also regulated by the redundant action of both genes. By contrast, an element of sub-functionalization is apparent in the C_4_ species maize and setaria, with the *GLK2* paralog expressed preferentially in bundle sheath cells. However, data from a number of different C_3_, C_4_ and intermediate species (Washburn *et al*. 2021) (Figure S1) invoke a more complex scenario because preferential expression of *GLK2* paralogs in bundle sheath cells is evident in species where decarboxylation reactions occur in the bundle sheath chloroplasts (NADP-ME type) but not in species where decarboxylation occurs in the mitochondria (NAD-ME type) or in the cytoplasm (PEP-CK type). This suggests that preferential localization of GLK2 in bundle sheath cells is associated with specific aspects of chloroplast function as opposed to general activation of photosynthesis.

The evolution of C_4_ required activation of photosynthesis and a significant increase in chloroplast volume in bundle sheath cells and as such, selection on regulatory regions of *GLK2* genes to enable high levels of activity in that cell type may have been a critical step. Notably, there is more within-species sequence variation in *GLK2* promoters of different rice and maize varieties than in the corresponding *GLK1* promoters (Figure S6), providing more opportunity for change through drift or selection. *GLK2* promoter sequences are also less conserved between species than corresponding *GLK1* sequences (Figure S7). ATAC-seq data from leaves shows that the proximal promoter and 5’UTR regions are the main sites of protein binding in *GLK* genes from maize, setaria and rice (Figure S4), with noticeably more DNA binding motifs related to light responsive elements in the 5’ UTR sequences of *GLK2* genes than in equivalent regions of *GLK1* genes. In maize, an enrichment of BPC motifs is seen in the *ZmG2* 5’UTR and the sequences are almost completely conserved across 1200 maize varieties. This level of sequence conservation in the 5’UTR is in stark contrast to the rest of the promoter region which is highly variable (Figure S6). Intriguingly, very few DNA binding motifs are found in either the *SvGLK1* or *SvGLK2* promoters (Figure S4) and no variation relevant to the domestication process was found when compared to *Setaria italica* (He *et al*. 2023). As in *ZmG2*, however, BPC motifs are present in both the *SvGLK2* and *OsGLK2* 5’UTR sequences but they are not found in any of the *GLK1* gene sequences. This observation suggests that the *GLK1* and *GLK2* 5’ UTR sequences diverged in the last common ancestor of maize, setaria and rice and thus prior to the evolution of C_4_. Because BPC binding proteins act to recruit polycomb complex proteins and thus mediate chromatin changes (Theune *et al*. 2019), this divergence may have enabled novel post-transcriptional regulation of *GLK*2 orthologs during the evolution of C_4_. Differences in the presence and position of upstream open reading frames (uORFs) within the 5’ UTR may have further facilitated de-repression of GLK2 translation (Zhang *et al*. 2020) in that putative uORFs in ZmGLK1 and SvGLK1 are situated less than 10 base pairs after the transcription start site (TSS) and thus if functional are predicted to be highly repressive, whereas the uORF in ZmGLK2 is over 200 bp downstream of the TSS and no uORFs are predicted in SvGLK2 (Figure S7). In rice, predicted uORFs are positioned 100 bp downstream of the TSS in both genes and may thus have no differential effect. Although requiring experimental verification, collectively these observations support the hypothesis that altered *GLK2* gene regulation was associated with the evolution of functional chloroplasts in bundle sheath cells of C_4_ grasses.

## METHODS

### Plant growth conditions

Maize and *S. viridis* seeds were germinated in hydrated vermiculite or on paper, respectively, and placed in an Aralab Fitoclima D1200 PLH incubator under 12 h light/12 h dark cycle at an intensity of 320 μmol photon m^−2^ s^−1^, with temperatures ranging from 31 °C during the day to 22 °C at night and humidity between 50-60 %. About 7-10 days after sowing (DAS), seedlings were transferred to pots containing John Innes no. 2 soil mixture and irrigated with water to keep the soil moist. Plants were kept under these conditions until sampled for experimental analysis.

For genetic crosses, maize plants were transferred to larger pots with the same soil mixture supplemented with Osmocote slow-release granules. Plants were grown in a greenhouse at the Department of Biology at the University of Oxford, UK with 16 h light/8 h dark cycle, daytime temperature 28 °C and night-time temperature 20 °C, with supplemental light provided when natural light levels were below 120 μmol photon m^−2^ s^−1^.

### Construct design and assembly

*ZmG2* and *ZmGLK1* orthologs were identified in *S. viridis* via phylogenetic reconstruction of the GARP gene family (see Figure S2). Guide RNAs (gRNA) to drive CRISPR/Cas9 site-directed mutagenesis were designed using the CRISPOR (Concordet and Haeussler 2018) online tool (http://crispor.tefor.net/) to achieve high editing efficiency for the target genes *ZmGLK1* (Zm00001d044785), *SvGLK1* (Sevir.4G123200) and *SvGLK2* (Sevir.5G003100), and to minimize off-targets. For maize, the designed gRNAs were sent to the Wisconsin Crop Innovation Centre (University of Wisconsin) to be cloned and transformed using their standard pipeline for maize transformation. For *S. viridis*, the gRNA sequences were synthesized as oligonucleotides with appropriate flanking sites for Golden Gate cloning (Table S1; JLF_Ox43 and JLF_Ox44 for *SvGLK1*; JLF_Ox51 and JLF_Ox52 for *SvGLK2*). Prior to cloning, the oligonucleotides were annealed and phosphorylated by incubating 10 μM of each with 1x Ligase Buffer (New England Biolabs) and 10 U of T4 Polynucleotide Kinase (New England Biolabs) in a final reaction volume of 15 μl at 37°C for 1.5 hours. After that period, 2 μl of 0.5 M NaCl was added, and the reaction was placed in a water bath heated to 95 °C for 2 min. The reaction was cooled to room temperature before being diluted 100-fold and cloned downstream of the *TaU6* promoter (module pJG310) using the Golden Gate one-step one-pot protocol (Engler *et al*. 2009) with an appropriate restriction enzyme (*Esp3I*). This module was then combined with another level 1 module containing *TaCas9* downstream of the maize ubiquitin promoter (pJG471) into the final acceptor and binary vector pTC278. Both modules and the vector were provided by Dr Dan Voytas (U. of Minnesota).

To overexpress *ZmGLK1* and *ZmG2*, the coding sequences of the maize genes were domesticated to avoid recognition sites for the type II restriction enzymes used in Golden Gate assembly: *BsaI*, *BpiI*, *Esp3I* and *DraIII*. The final sequence was synthesized as a level 0 module with the appropriate SC flanking sites (Engler and Marillonnet 2013). *ZmGLK1* and *ZmG2* level 0 modules (EC17054 and EC17056) were cloned downstream of the maize ubiquitin promoter (*ZmUBI*_pro_, EC15455) and upstream of a *nos* terminator (tNOS, EC41421) in level 1 backbone position forward 2 using the Golden Gate one-step one-pot protocol (Engler *et al*. 2009). The previously described *hygromycin phosphotransferase* (HygR) coding sequence cloned downstream of the rice actin promoter (*OsACT*_pro_) was used as the selectable marker (Vlad *et al*. 2019). Level 1 modules were assembled into the binary vector pAGM4723 to obtain the constructs EC17808 and EC17810 (hereafter referred to as 17808 and 17810, respectively) depicted in Figure S5B and S5C.

### Plant transformation

*S. viridis* accessions A10 and ME034V were used to obtain CRISPR/Cas9-driven mutations for *SvGLK1* and *SvGLK2*, respectively. *Agrobacterium tumefaciens* strain AGL1 harbouring the construct of interest was co-cultivated with callus generated from seeds of *S. viridis*. Callus induction, transformation and seedling regeneration were performed as described by VanEck and collaborators (Van Eck and Startwood 2015). Complementation lines were obtained by transforming constructs 17808 and 17810 into callus generated from a population of seeds that segregated for the gene-edited *Svglk2* allele but no longer segregated the original t-DNA which had been recombined out. The presence of t-DNA in regenerated seedlings was confirmed via duplex PCR (Table S1; JLF_Ox12 and JLF_Ox13 amplify the HygR gene; JLF_Ox14 and JLF_Ox15 amplify an endogenous gene used as a DNA quality control). The occurrence of gene editing by CRISPR/Cas9 was verified by PCR amplification of the region flanking the expected mutation site (Table S1; JLF_Ox45 and JLF_Ox46 for *SvGLK1*; JLF_Ox53 and JLF_Ox54 for *SvGLK2*) followed by endonuclease digestion with *MmeI* or *Tsp45I* for *SvGLK1* and *SvGLK2*, respectively, following the manufacturers’ guidelines.

*Z. mays* Hi-II was transformed by the Wisconsin Crop Innovation Centre (University of Wisconsin) according to their transformation pipeline. The presence of the t-DNA was assessed in T1 lines by PCR (Table S1; JLF_Ox178 and JLF_Ox179) and *Zmglk1* mutants were identified by PCR amplification of the region flanking the expected mutation site (Table S1; JLF_Ox39 and JLF_Ox175) followed by endonuclease digestion with *HinfI* according to the manufacturer’s instructions.

For all lines, mutated sequences were verified by Sanger sequencing (Source BioScience, UK). Experimental analyses were performed on homozygous plants in the T2 or subsequent generations.

### Confocal imaging

The mid-portion of fully expanded leaf 7 from 21 DAS *S. viridis* plants was fixed in 4 % paraformaldehyde/0.2 % glutaraldehyde in 25 mM phosphate buffer pH 7.2. After fixation, leaves were stored in 0.2 M Na-EDTA pH 9.0. Free-hand cross sections were imaged using a Leica SP9 confocal microscope with an Argon laser set to excite chlorophyll at 663 nm and to emit from 650-800 nm.

### Ultrastructure analysis via transmission electron microscopy

Sample fixation and preparation were performed by the Electron Microscopy Facility team at the Sir Willian Dunn School of Pathology (University of Oxford). Leaf discs of 2 mm diameter were collected from leaf 5 of 40 DAS maize or 25 DAS *S. viridis* plants and placed into 3 % glutaraldehyde in 0.06 M Sorenson’s phosphate buffer pH 7.2. Samples were transferred to a Leica AMW microwave tissue processor for fixation, osmification (with 1 % OsO_4_), dehydration with an acetone series, and resin infiltration (LRW Hard). The samples were then removed from the microwave and transferred to fresh resin for four days, renewing the resin twice a day. The embedded samples were transferred into gelatin capsules and filled with resin for polymerization at 60 °C for about 65 h. Sections of 150 nm were cut using a Leica UC7 ultramicrotome, collected onto formvar-coated copper slot grids and post-stained with lead citrate. Images were acquired on a Thermo Fischer Tecnai T12 TEM with a Gatan OneView CMOS camera.

Thylakoid occupancy of chloroplasts was quantified using ImageJ. The thylakoid area was measured by adjusting the image threshold to create a binary mask on which only the thylakoid membrane was apparent. This area value was divided by the chloroplast planar area to give the thylakoid occupancy level.

### Chlorophyll extraction and quantification

Leaf discs were harvested from fully expanded leaf 5 at 30 DAS for maize and 21 DAS for *S. viridis*, and immediately frozen in liquid nitrogen. For total chlorophyll extraction, finely powdered tissue was resuspended in 80 % acetone buffered in 100 mM Tris-HCl pH 8.0 (Chazaux *et al*. 2022). Following a centrifugation step, the absorbance of each sample was read in technical triplicates in a FLUOstar Omega spectrometer (BMG Labtech) at 646 and 663 nm. Background absorbance was measured at 750 nm. Absorbance values recorded at 750 nm were subtracted from values recorded at 646 and 663 nm for normalization purposes. Technical replicates were averaged for each sample. Concentration of chlorophylls a + b were calculated in micrograms (μg) per leaf area using the following the equation: (19.54 × Abs_646_ + 8.29 × Abs_663_) × dilution factor (Porra *et al*. 1989).

### Fluorescence quantification of PSII

Photosystem II (PSII) fluorescence was measured using a LiCOR Li-6800 portable photosynthesis system. LiCOR chamber conditions were set to a flow intensity of 500 umol s^−1^, delta pressure of 0.2 kPa, relative air humidity of 50 %, reference concentration of CO_2_ at 400 umol mol^−1^ and temperature exchange at 25 °C. For quantification of PSII operating efficiency (PhiPSII), plants were adapted to light under the conditions described above for at least 1.5 h and actinic light intensity on the LiCOR chamber was set to 1000 μmol photon m^−2^ s^−1^. After clamping, each leaf was adapted for a few minutes until stabilization was reached before logging the fluorescence value. Quantification of maximum quantum efficiency of PSII photochemistry (Fv/Fm) was performed on plants that were dark-adapted for at least 16 h. LiCOR actinic lights were off. In both cases, the fluorescence action log was set to FoFm (dark), FsFm’ (light), rectangular flash type with red target at 8000 μmol photon m^−2^ s^−1^, duration of 1000 ms, output rate of 100 Hz and margin of 5 points.

### Gene expression analysis via qPCR

The mid-portion of fully expanded leaf 5 from at least three biological replicates of 30 DAS maize or 21 DAS *S. viridis* were harvested and immediately frozen in liquid nitrogen. Tissue was ground prior to total RNA extraction using a RNeasy Plant Mini Kit (Qiagen) according to the manufacturer’s instructions. To avoid any DNA contamination, samples were treated with TURBO DNA-free DNase (Thermo Fischer Scientific) according to the manufacturer’s guidelines. For first strand cDNA synthesis, 100 ng of total RNA was added to 1 μl of Maxima Enzyme Mix (Thermo Fischer Scientific) and 1x Reaction Mix to a final volume of 10 μL. The mixture was incubated at 25 °C for 10 min, followed by 50 °C for 15 min. Enzyme inactivation was carried out at 85 °C for 5 min.

Amplification reactions were performed in a StepOnePlus Real-Time PCR System (Thermo Fischer Scientific) using SYBRGreen to monitor dsDNA synthesis. The reaction mixtures contained 5 μL of diluted cDNA (1:25), 0.2 μM of each primer and 10 μL of SYBR™ Green PCR Master Mix (Thermo Fischer Scientific) in a total volume of 10 μL. The reaction cycles began with a 5 min denaturation step at 94 °C, followed by 40 amplification cycles of 15 s at 94 °C, 10 s at 60 °C, 15 s at 72 °C. After each cycle, the fluorescence was measured at 60 °C for 35 s. The melting curve was produced by a cycle of 95 °C for 15 s, 60 °C for 1 min, 95 °C for 30 s and finally 60 °C for 15 s. Every qPCR reaction was repeated three times to make technical replicates.

Transcript levels for multiple target genes were measured with primers designed to specifically amplify the gene of interest (Table S1). Two reference genes were used to normalize gene expression for maize (Lin *et al*. 2014) (Table S1; ZmCYP_F and ZmCYP_R, ZmEF1α_F and ZmEF1α_R) and *S. viridis* (Lambret-Frotté *et al*. 2015) (Table S1; JLF_Ox26 and JLF_Ox27, JLF_Ox28 and JLF_Ox29). Exported *Rn* values were used to calculate amplification efficiency for each primer pair and the Cq values for each qPCR reaction using the ‘*qpcR*’ package from R (Ritz and Spiess 2008). Normalized relative expression of the target genes was calculated using the R package ‘*EasyqpcR’* (https://www.bioconductor.org/packages//2.12/bioc/html/EasyqpcR.html), according to the ΔΔCq model (Pfaffl 2001) based on the previously calculated Cq and amplification efficiencies.

### Statistical analysis

Datasets were tested for Normal distributions using the Shapiro-Wilk test (Shapiro and Wilk 1965). For thylakoid coverage, chlorophyll concentration and PSII fluorescence, variance was calculated using a one-way ANOVA followed by a pairwise comparison analysis with Tukey’s HSD test (Tukey 1949). For relative gene expression, statistical significance was calculated based on a Student’s t-test, comparing the mean variance of each mutant line with their respective controls. All statistical analyses were performed using *scipy* v.1.5.0 library and plots were prepared using *matplotlib* v3.5.1 library; both from Python v.3.7.6, using Visual Studio Code as a code editor.

## LIST OF SUPPLEMENTARY MATERIAL

Table S1. Oligonucleotides used for cloning, genotyping and qPCR.

Table S2. Data and statistical analyses supporting Figures 1-6.

Figure S1. *GLK* transcript accumulation in bundle sheath and mesophyll cells.

Figure S2. Phenotypic characterization of three independent *Zmglk1* mutant lines.

Figure S3. Comparison of *GLK* sequences.

Figure S4. DNA binding motifs in regulatory regions of *GLK* genes.

Figure S5. Phenotypic characterization of *Svglk2* mutant lines complemented with *ZmGLK1* or *ZmG2*.

Figure S6. Sequence variability in regulatory regions of *GLK* genes.

Figure S7. Alignment of *GLK* gene promoter and 5’ UTR regions.

## AUTHOR CONTRIBUTIONS

J.L-F and J.A.L conceived and designed the experiments; G.S carried out setaria transformation, plant cultivation and genotyping; J.L-F carried out all of the remaining experiments and analysed the data. J.L-F and J.A.L wrote the manuscript.

## ACKNOWLEDGEMENTS

The authors thank Dan Voytas for making available the CRISPR/Cas9 cloning vectors for gene editing in setaria; Raman Dhaliwal, Charlotte Melia and Errin Jonson for processing and imaging the TEM samples; Ray Collier, Mike Pettersen and the Wisconsin Crop Innovation Centre team for maize transformation; John Baker for plant photography; Roxaana Clayton, Julie Bull and Lizzie Jamison for technical support; Emma Raven for assisting with initial characterization of mutant lines; Jacob Washburn for providing TPM data for Figure S1B; Sophie Johnson, Chiara Perico, Daniela Vlad, Sovanna Tan, Thomas Hughes and Maricris Zaidem for discussion throughout the experimental work and during manuscript preparation.

## SUPPLEMENTARY FIGURE LEGENDS

**Figure S1. *GLK* transcript accumulation in bundle sheath and mesophyll cells. A)** Gene expression in reads per kilobase million (RPKM) in mesophyll (MC) and bundle sheath cells (BSC) of setaria (*SvPEPC*, *SvNADP*-*ME*, *SvGLK1*, *SvGLK2*) and maize (*ZmPEPC*, *ZmNADP*-*ME*, *ZmGLK1*, *ZmG2*). Maize data were published by Li et al. (2010) and Chang et al. (2012), and setaria data were published by John et al. (2014). *PEPC* and *NADP-ME* transcripts accumulate specifically in mesophyll and bundle sheath cells respectively, and thus act as markers for cross contamination of RNA samples between cell-types. As such, mesophyll contamination of bundle sheath transcriptomes was 22% (Li et al), 4.5% (Chang et al) and 3.5% (John et al), whereas bundle sheath contamination of mesophyll transcriptomes was 10.7% (Li et al), 7% (Chang et al) and 6.6% (John et al). Disregarding the Li et al (2010) data because of the high level of cross contamination, *GLK1* transcript levels were higher in mesophyll than bundle sheath cells by 8-fold in maize and 21-fold in setaria, whereas *GLK2* transcript levels were higher in bundle sheath than mesophyll cells by 4.8-fold in maize and 3.2-fold in setaria. **B)** Log2 fold change between bundle sheath and mesophyll transcripts for *S. viridis* (John et al. 2014), *Z. mays* (Li et al. 2010, Chang et al. 2012), and three different species of Paniceae grasses (*Urochloa fusca*, *Panicum hallii* and *Digitaria californica*) (Washburn et al. 2022). All species carry out C_4_ photosynthesis, but each uses a different decarboxylation pathway in the bundle sheath cells. *PEPC* was used as a mesophyll cell marker in each case and genes encoding the relevant decarboxylation enzyme (i.e. *NADP-ME, PCK* and *NAD-ME*) were used as bundle sheath cell markers.

**Figure S2. Phenotypic characterization of three independent *Zmglk1* mutant lines. A-D)** Whole plant phenotype 30 days after sowing. Scale bars = 10 cm. **E-H)** Transmission electron microscopy (TEM) images showing mesophyll chloroplast ultrastructure. Scale = 2 μm, 2 μm, 5 μm and 2 μm, respectively. **I-L)** TEM images showing bundle sheath chloroplast ultrastructure. Scale bars = 2 μm, 5 μm, 10 μm and 2 μm, respectively.

**Figure S3. Comparison of GLK sequences**. **A)** Phylogenetic reconstruction of the *GLK* gene family. Protein sequences of the extensively characterised GLK members ZmGLK1, ZmG2, OsGLK1 and OsGLK2 were used to create a HMMER profile that was used as a query to search for homologs in whole protein databases of 11 species (*Zea mays, Setaria viridis, S. italica, Panicum virgatum, Sorghum bicolor, Oryza sativa, Brachypodium distachyon, Vitis vinifera, Arabidopsis thaliana, Marchantia polymorpha* and *Physcomitrium patens*). The presence of the MYB domain in those sequences was confirmed using InterproScan, and only the sequences containing the conserved AREAEAA domain characteristic of the GLK subfamily were selected. The resulting 25 amino acid sequences were aligned using Clustal Omega. Phylogenetic reconstruction was performed using Bayesian inference on Mr. Bayes using Poisson with fixed substitution rates as a model. Two independent MCMC runs with 1,000,000 generations were performed and no further improvement of log likelihood of prior probabilities was observed. The tree was visualized on FigTree. Clade support on each node was assessed using Bayesian posterior probabilities. Members of the GLK1 subgroup are depicted in green and GLK2 in blue. **B)** Amino acid sequence alignment showing conservation between ZmGLK1, SvGLK1, ZmG2 and SvGLK2. The diagonal linesl show the HLH associated with MYB or MYB-related proteins; the AREAEAA conserved site is depicted in horizontal lines and the last exon that contains the GCT-box domain is depicted in vertical lines.

**Figure S4. DNA binding motifs in regulatory regions of *GLK* genes**. **A)** Schematics of maize, setaria and rice *GLK* genes showing binding sites revealed from ATAC assays with leaf RNA. Data was retrieved from https://epigenome.genetics.uga.edu/PlantEpigenome/index.html. The yellow shading highlights the 800 bp upstream of the transcription start site plus the 5’UTR regions. **B)** Table showing the number of DNA binding motifs found in regulatory regions of each gene. Plant non-redundant motifs (JASPAR) was used to screen the above-mentioned regions of each gene, using the FIMO tool with a p value of <0.0001. The photosynthesis-related/light responsive motifs (BPC, ABI3-like, GT-1, GT-4, CNA, MYB-B, STZ, bZIP/bHLH(G-box), GATA, MYB-like/I-box, TCP, TGA and WRKY) were as described by Sing et al. (2023) and rice and maize GLK-specific binding sites were as described in Tu et al. (2022). **C, D)** Schematics of 800 bp upstream of the transcription start site plus the 5’UTR regions for each gene, showing the position of all predicted DNA binding motifs (C) and of known light-responsive motifs (D). The 5’ UTR is shown in white and promoter sequence in grey.

**Figure S5. Phenotypic characterization of *Svglk2* mutant lines complemented with *ZmGLK1* or *ZmG2***. **A)** Schematic of the gene edited *Svglk2* mutant allele. **B-C)** Schematic of the constructs used to express *ZmGLK1* (B) or *ZmG2* (C) in the *Svglk2* background. *HygR* depicts the hygromycin phosphotransferase gene and *OsACT_pro_* and *ZmUBI_pro_* represent the constitutive rice actin and maize ubiquitin promoters, respectively. LB and RB refer to left and right borders. **D-M)** Transmission electron microscopy images showing mesophyll (MC) and bundle sheath (BSC) cell chloroplast ultrastructure in the mutant line (D, E) and in lines overexpressing *ZmGLK1* (F-I) or *ZmG2* (J-M) in the mutant background. Scale bar sizes are indicated on each image. D & E are the same images as Figure 3G & 3H). **N-R)** Confocal images of leaf cross sections from the mutant (N) and from lines overexpressing *ZmGLK1* (O, P) or *ZmG2* (Q, R) showing morphology of chloroplasts. Bundle sheath cells are outlined in white. Scale bars = 50 μm. Yellow arrows indicate ectopic chloroplast formation in vascular cells

**Figure S6. Sequence variability in regulatory regions of *GLK* genes.** The SNPs from 3025 variants of rice (SNP seek, IRRI) and 1210 variants of maize (Maize SNPDB) were used to calculate the frequency of each nucleotide for each SNP position. The nucleotide with higher prevalence in each position is plotted on the graphs for *OsGLK1*, *OsGLK2, ZmGLK1* and *ZmG2*. The 2Kb upstream of the transcription start site is shown in grey, the 5’UTR in white and the first exon in black.

**Figure S7. Alignment of *GLK* gene promoter and 5’ UTR regions**. **A, B)** *GLK1* (B) and *GLK2 (*A) sequences (800 bp upstream of the transcription start site (TSS) plus the 5’ UTR) aligned using MAFFT (Katoh *et al*. 2019) and visualized using JALview (Waterhouse *et al*. 2009). Conserved bases are indicated by blue shading, the predicted TATA box by the red squares, actual (rice - Murray et al. 2022) or predicted TSS by orange squares and putative uORFs by yellow squares. **C)** Amino acid sequences of the predicted uORFs in (A) & (B).

## REFERENCES

Bravo-Garcia, A., Yasumura, Y., and Langdale, J.A. (2009) Specialization of the *Golden2-like* regulatory pathway during land plant evolution. New Phytologist 183, 133–141.

Chang, Y.-M., Liu, W.-Y., Shih, A.C.-C., Shen, M.-N., Lu, C.-H., Lu, M.-Y.J., Yang, H.-W., Wang, T.-Y., Chen, S.C.-C., Chen, S.M., Li, W.-H., and Ku, M.S.B. (2012) Characterizing Regulatory and Functional Differentiation between Maize Mesophyll and Bundle Sheath Cells by Transcriptomic Analysis. Plant Physiology 160, 165–177.

Chazaux, M., Schiphorst, C., Lazzari, G., and Caffarri, S. (2022) Precise estimation of chlorophyll a, b and carotenoid content by deconvolution of the absorption spectrum and new simultaneous equations for Chl determination. Plant Journal 109, 1630–1648.

Chiang, Y.-H., Zubo, Y.O., Tapken, W., Kim, H.J., Lavanway, A.M., Howard, L., Pilon, M., Kieber, J.J., and Schaller, G.E. (2012). Functional Characterization of the GATA Transcription Factors GNC and CGA1 Reveals Their Key Role in Chloroplast Development, Growth, and Division in Arabidopsis. Plant Physiology 160, 332–348.

Christin, P.-A. and Osborne, C.P. (2013) The recurrent assembly of C_4_ photosynthesis, an evolutionary tale. Photosynthesis Research 117, 163–75.

Concordet, J.-P. and Haeussler, M. (2018) CRISPOR: intuitive guide selection for CRISPR/Cas9 genome editing experiments and screens. Nucleic Acids Research 46, W242–W245.

Cribb, L., Hall, L., and Langdale, J.A. (2001) Four Mutant Alleles Elucidate the Role of the G2 Protein in the Development of C_4_ and C_3_ Photosynthesizing Maize Tissues. Genetics 159 787–797.

Van Eck, J. and Startwood, K. (2015) Setaria viridis. In: K. Wang (Ed.). Agrobacterium Protocols. Springer, 417.

Engler, C., Gruetzner, R., Kandzia, R., and Marillonnet, S. (2009) Golden gate shuffling: a one-pot DNA shuffling method based on type IIs restriction enzymes. PloS ONE 4, e5553.

Engler, C. and Marillonnet, S. (2013) Combinatorial DNA assembly using golden gate cloning. Methods in Molecular Biology 1073, 141–156.

Fitter, D.W., Martin, D.J., Copley, M.J., Scotland, R.W., and Langdale, J.A. (2002) *GLK* gene pairs regulate chloroplast development in diverse plant species. Plant Journal 31, 713–727.

Gallaher, T.J., Peterson, P.M., Soreng, R.J., Zuloaga, F.O., Li, D.-Z., Clark, L.G., Tyrrell, C.D., Welker, C.A.D., Kellogg, E.A., and Teisher, J.K. (2022) Grasses through space and time: An overview of the biogeographical and macroevolutionary history of Poaceae. Journal of Systematics and Evolution 60, 522–569.

Hall, L.N., Rossini, L., Cribb, L., and Langdale, J.A. (1998) *GOLDEN 2*: A novel transcriptional regulator of cellular differentiation in the maize leaf. Plant Cell 10, 925–936.

He, Q., Tang, S., Zhi, H., Chen, J., Zhang, J., Liang, H., Alam, O., Li, H., Zhang, H., Xing, L., Li, X., Zhang, W., Wang, H., Shi, J., Du, H., Wu, H., Wang, L., Yang, P., Xing, L., Yan, H., Song, Z., Liu, J., Wang, H., Tian, X., Qiao, Z., Feng, G., Guo, R., Zhu, W., Ren, Y., Hao, H., Li, M., Zhang, A., Guo, E., Yan, F., Li, Q., Liu, Y., Tian, B., Zhao, X., Jia, R., Feng, B., Zhang, J., Wei, J., Lai, J., Jia, G., Purugganan, M., and Diao, X. (2023) A graph-based genome and pan-genome variation of the model plant *Setaria*. Nature Genetics 55, 1232–1242

Hua, L., Stevenson, S.R., Reyna-Llorens, I., Xiong, H., Kopriva, S., and Hibberd, J.M. (2021) The bundle sheath of rice is conditioned to play an active role in water transport as well as sulfur assimilation and jasmonic acid synthesis. Plant Journal 107, 268–286.

Hudson, D., Guevara, D.R., Hand, A.J., Xu, Z., Hao, L., Chen, X., Zhu, T., Bi, Y.-M., and Rothstein, S.J. (2013) Rice Cytokinin GATA Transcription Factor1 Regulates Chloroplast Development and Plant Architecture. Plant Physiology 162, 132–144.

John, C.R., Smith-Unna, R.D., Woodfield, H., Covshoff, S., and Hibberd, J.M. (2014) Evolutionary convergence of cell-specific gene expression in independent lineages of C_4_ grasses. Plant Physiology 165, 62–75.

Katoh, K., Rozewicki, J., and Yamada, K.D. (2019) MAFFT online service: multiple sequence alignment, interactive sequence choice and visualization. Briefings in Bioinformatics 20, 1160–1166.

Kenrick, P. and Crane, P.R. (1997) The origin and early evolution of plants on land. Nature 389, 33–39.

Kobayashi, K., Baba, S., Obayashi, T., Sato, M., Toyooka, K., Keränen, M., Aro, E.M., Fukaki, H., Ohta, H., Sugimoto, K., and Masuda, T. (2012) Regulation of root greening by light and auxin/cytokinin signaling in Arabidopsis. Plant Cell, 24, 1081–1095.

Lambret-Frotté, J., Almeida, L.C.S. De, Moura, S.M. De, and Souza, F.L.F. (2015) Validating Internal Control Genes for the Accurate Normalization of qPCR Expression Analysis of the Novel Model Plant *Setaria viridis*. PloS One 10, e0135006.

Langdale, J.A. (2011) C_4_ cycles: past, present, and future research on C_4_ photosynthesis. The Plant Cell 23, 3879–92.

Langdale, J.A. and Kidner, C.A. (1994) *bundle sheath defective*, a mutation that disrupts cellular differentiation in maize leaves. Development 120, 673–681.

Li, P. and Brutnell, T.P. (2011) *Setaria viridis* and *Setaria italica*, model genetic systems for the Panicoid grasses. Journal of Experimental Botany 62, 3031–7.

Li, P., Ponnala, L., Gandotra, N., Wang, L., Si, Y., Tausta, S.L., Kebrom, T.H., Provart, N., Patel, R., Myers, C.R., Reidel, E.J., Turgeon, R., Liu, P., Sun, Q., Nelson, T., and Brutnell, T.P. (2010) The developmental dynamics of the maize leaf transcriptome. Nature Genetics 42, 1060–7.

Li, X., Wang, P., Li, J., Wei, S., Yan, Y., Yang, J., Zhao, M., Langdale, J.A., and Zhou, W. (2020) Maize *GOLDEN2-LIKE* genes enhance biomass and grain yields in rice by improving photosynthesis and reducing photoinhibition. Communications Biology 3, 151.

Lin, Y., Zhang, C., Lan, H., Gao, S., Liu, H., Liu, J., Cao, M., Pan, G., Rong, T., and Zhang, S. (2014) Validation of potential reference genes for qPCR in maize across abiotic stresses, hormone treatments, and tissue types. PloS One 9, e95445.

Marshall, D.M., Muhaidat, R., Brown, N.J., Liu, Z., Stanley, S., Griffiths, H., Sage, R.F., and Hibberd, J.M. (2007) Cleome, a genus closely related to Arabidopsis, contains species spanning a developmental progression from C_3_ to C_4_ photosynthesis. Plant Journal 51, 886–896.

Murray, A., Mendieta, J.P., Vollmers, C., and Schmitz, R.J. (2022) Simple and accurate transcriptional start site identification using Smar2C2 and examination of conserved promoter features. The Plant Journal 112, 583–596.

Nguyen, C. V., Vrebalov, J.T., Gapper, N.E., Zheng, Y., Zhong, S., Fei, Z., and Giovannoni, J.J. (2014) Tomato *GOLDEN2-LIKE* transcription factors reveal molecular gradients that function during fruit development and ripening. Plant Cell 26, 585–601.

Nickelsen, J. and Rengstl, B. (2013) Photosystem II Assembly: From Cyanobacteria to Plants. Annual Review of Plant Biology 64, 609–635.

Pfaffl, M.W. (2001) A new mathematical model for relative quantification in real-time RT-PCR. Nucleic Acids Research 29, e45.

Porra, R.J., Thompson, W.A., and Kriedemann, P.E. (1989) Determination of accurate extinction coefficients and simultaneous equations for assaying chlorophylls a and b extracted with four different solvents: verification of the concentration of chlorophyll standards by atomic absorption spectroscopy. Biochimica et Biophysica Acta (BBA) - Bioenergetics 975, 384–394.

Riechmann, J.L., Heard, J., Martin, G., Reuber, L., Jiang, C., Keddie, J., Adam, L., Pineda, O., Ratcliffe, O.J., Samaha, R.R., Creelman, R., Pilgrim, M., Broun, P., Zhang, J.Z., Ghandehari, D., Sherman, B.K., and Yu, G. (2000) Arabidopsis transcription factors: genome-wide comparative analysis among eukaryotes. Science 290, 2105–2110.

Ritz, C. and Spiess, A.-N. (2008) qpcR: an R package for sigmoidal model selection in quantitative real-time polymerase chain reaction analysis. Bioinformatics 24, 1549–1551.

Rossini, L., Cribb, L., Martin, D.J., and Langdale, J.A. (2001) The maize *Golden2* gene defines a novel class of transcriptional regulators in plants. Plant Cell 13, 1231–1244.

Safi, A., Medici, A., Szponarski, W., Ruffel, S., Lacombe, B., and Krouk, G. (2017) The world according to GARP transcription factors. Current Opinion in Plant Biology 39, 159–167.

Sagan, L. (1967) On the origin of mitosing cells. Journal of Theoretical Biology 14, 225–IN6.

Sage, R.F., Christin, P.-A., and Edwards, E.J. (2011) The C_4_ plant lineages of planet Earth. Journal of Experimental Botany 62, 3155–69.

Shapiro, S.S. and Wilk, M.B. (1965) An analysis of variance test for normality (complete samples). Biometrika 52, 591–611.

Tausta, L.S., Li, P., Si, Y., Gandotra, N., Liu, P., Sun, Q., Brutnell, T.P., and Nelson, T. (2014) Developmental dynamics of Kranz cell transcriptional specificity in maize leaf reveals early onset of C_4_-related processes. Journal of Experimental Botany 65, 3543–3555.

Theune, M.L., Bloss, U., Brand, L.H., Ladwig, F., and Wanke, D. (2019) Phylogenetic analyses and GAGA-motif binding studies of BBR/BPC proteins lend to clues in GAGA-MOTIF recognition and a regulatory role in brassinosteroid signaling. Frontiers in Plant Science 10, e00466.

Tu, X., Ren, S., Shen, W., Li, J., Li, Y., Li, C., Li, Y., Zong, Z., Xie, W., Grierson, D., Fei, Z., Giovannoni, J., Li, P., and Zhong, S. (2022) Limited conservation in cross-species comparison of GLK transcription factor binding suggested wide-spread cistrome divergence. Nature Communications 13, 7632.

Tukey, J.W. (1949) Comparing Individual Means in the Analysis of Variance. Biometrics 5, 99–114.

Vlad, D., Abu-Jamous, B., Wang, P., and Langdale, J.A. (2019) A modular steroid-inducible gene expression system for use in rice. BMC Plant Biology 19, 426.

Wang, L., Czedik-Eysenberg, A., Mertz, R. a, Si, Y., Tohge, T., Nunes-Nesi, A., Arrivault, S., Dedow, L.K., Bryant, D.W., Zhou, W., Xu, J., Weissmann, S., Studer, A., Li, P., Zhang, C., LaRue, T., Shao, Y., Ding, Z., Sun, Q., Patel, R. V, Turgeon, R., Zhu, X., Provart, N.J., Mockler, T.C., Fernie, A.R., Stitt, M., Liu, P., and Brutnell, T.P. (2014) Comparative analyses of C_4_ and C_3_ photosynthesis in developing leaves of maize and rice. Nature Biotechnology, 32, 1158–65.

Wang, P., Fouracre, J., Kelly, S., Karki, S., Gowik, U., Aubry, S., Shaw, M.K., Westhoff, P., Slamet-Loedin, I.H., Quick, W.P., Hibberd, J.M., and Langdale, J.A. (2013) Evolution of *GOLDEN2-LIKE* gene function in C_3_ and C_4_ plants. Planta 237, 481–495.

Washburn, J.D., Strable, J., Dickinson, P., Kothapalli, S.S., Brose, J.M., Covshoff, S., Conant, G.C., Hibberd, J.M., and Pires, J.C. (2021) Distinct C_4_ sub-types and C_3_ bundle sheath isolation in the Paniceae grasses. Plant Direct 5, 1–14.

Waterhouse, A.M., Procter, J.B., Martin, D.M.A., Clamp, M., and Barton, G.J. (2009) Jalview Version 2— a multiple sequence alignment editor and analysis workbench. Bioinformatics 25, 1189–1191.

Waters, M.T. and Langdale, J.A. (2009) The making of a chloroplast. EMBO J. 28, 2861–2873.

Waters, M.T., Moylan, E.C., and Langdale, J.A. (2008) GLK transcription factors regulate chloroplast development in a cell-autonomous manner. Plant Journal 56, 432–444.

Waters, M.T., Wang, P., Korkaric, M., Capper, R.G., Saunders, N.J., and Langdale, J.A. (2009) GLK Transcription Factors Coordinate Expression of the Photosynthetic Apparatus in Arabidopsis. The Plant Cell 21, 1109–1128.

Wickett, N.J., Mirarab, S., Nguyen, N., Warnow, T., Carpenter, E., Matasci, N., Ayyampalayam, S., Barker, M.S., Burleigh, J.G., Gitzendanner, M.A., Ruhfel, B.R., Wafula, E., Der, J.P., Graham, S.W., Mathews, S., Melkonian, M., Soltis, D.E., Soltis, P.S., Miles, N.W., Rothfels, C.J., Pokorny, L., Shaw, A.J., DeGironimo, L., Stevenson, D.W., Surek, B., Villarreal, J.C., Roure, B., Philippe, H., dePamphilis, C.W., Chen, T., Deyholos, M.K., Baucom, R.S., Kutchan, T.M., Augustin, M.M., Wang, J., Zhang, Y., Tian, Z., Yan, Z., Wu, X., Sun, X., Wong, G.K.-S., and Leebens-Mack, J. (2014) Phylotranscriptomic analysis of the origin and early diversification of land plants. Proceedings of the National Academy of Sciences 111, E4859–E4868.

Woo, K.C., Anderson, J.M., Boardman, N.K., Downton, W.J.S., Osmond, C.B., and Thorne, S.W. (1970) Deficient Photosystem II in Agranal Bundle Sheath Chloroplasts of C_4_ Plants. Proceedings of the National Academy of Sciences 67, 18–25.

Yasumura, Y., Moylan, E.C., and Langdale, J.A. (2005) A Conserved Transcription Factor Mediates Nuclear Control of Organelle Biogenesis in Anciently Diverged Land Plants. Plant Cell 17, 1894– 1907.

Zhang, T., Wu, A., Yue, Y., and Zhao, Y. (2020) uORFs: Important cis-regulatory elements in plants. International Journal of Molecular Sciences 21, 1–4.

Zubo, Y.O., Blakley, I.C., Franco-Zorrilla, J.M., Yamburenko, M. V, Solano, R., Kieber, J.J., Loraine, A.E., and Schaller, G.E. (2018) Coordination of Chloroplast Development through the Action of the GNC and GLK Transcription Factor Families. Plant Physiology 178, 130–147.

