## Supplementary Information for "*GOLDEN2-like1* is sufficient but not necessary for chloroplast biogenesis in mesophyll cells of C_4_ grasses"

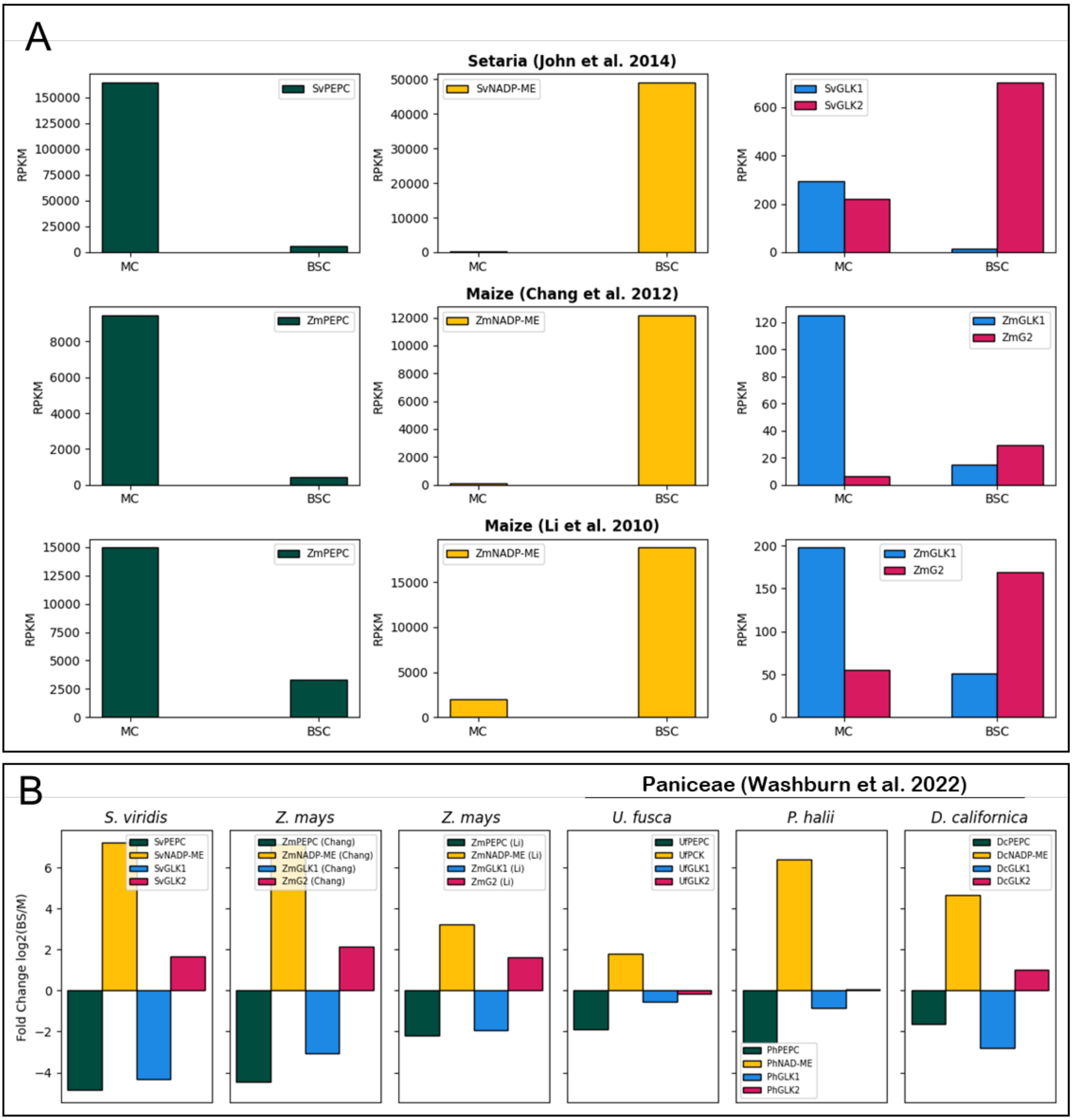


**Figure S1. *GLK* transcript accumulation in bundle sheath and mesophyll cells. A)** Gene expression in reads per kilobase million (RPKM) in mesophyll (MC) and bundle sheath cells (BSC) of setaria (*SvPEPC*, *SvNADP-ME*, *SvGLK1*, *SvGLK2*) and maize (*ZmPEPC*, *ZmNADP-ME*, *ZmGLK1*, *ZmG2*). Maize data were published by Li et al. (2010) and Chang et al. (2012), and setaria data were published by John et al. (2014). *PEPC* and *NADP-ME* transcripts accumulate specifically in mesophyll and bundle sheath cells respectively, and thus act as markers for cross contamination of RNA samples between cell-types. As such, mesophyll contamination of bundle sheath transcriptomes was 22% (Li et al), 4.5% (Chang et al) and 3.5% (John et al), whereas bundle sheath contamination of mesophyll transcriptomes was 10.7% (Li et al), 7% (Chang et al) and 6.6% (John et al). Disregarding the Li et al (2010) data because of the high level of cross contamination, *GLK1* transcript levels were higher in mesophyll than bundle sheath cells by 8-fold in maize and 21-fold in setaria, whereas *GLK2* transcript levels were higher in bundle sheath than mesophyll cells by 4.8-fold in maize and 3.2-fold in setaria. **B)** Log2 fold change between bundle sheath and mesophyll transcripts for *S. viridis* (John et al. 2014), *Z. mays* (Li et al. 2010, Chang et al. 2012), and three different species of Paniceae grasses (*Urochloa fusca*, *Panicum hallii* and *Digitaria californica*) (Washburn et al. 2022). All species carry out $C_4$ photosynthesis, but each uses a different decarboxylation pathway in the bundle sheath cells. *PEPC* was used as a mesophyll cell marker in each case and genes encoding the relevant decarboxylation enzyme (i.e. *NADP-ME, PCK* and *NAD-ME*) were used as bundle sheath cell markers.

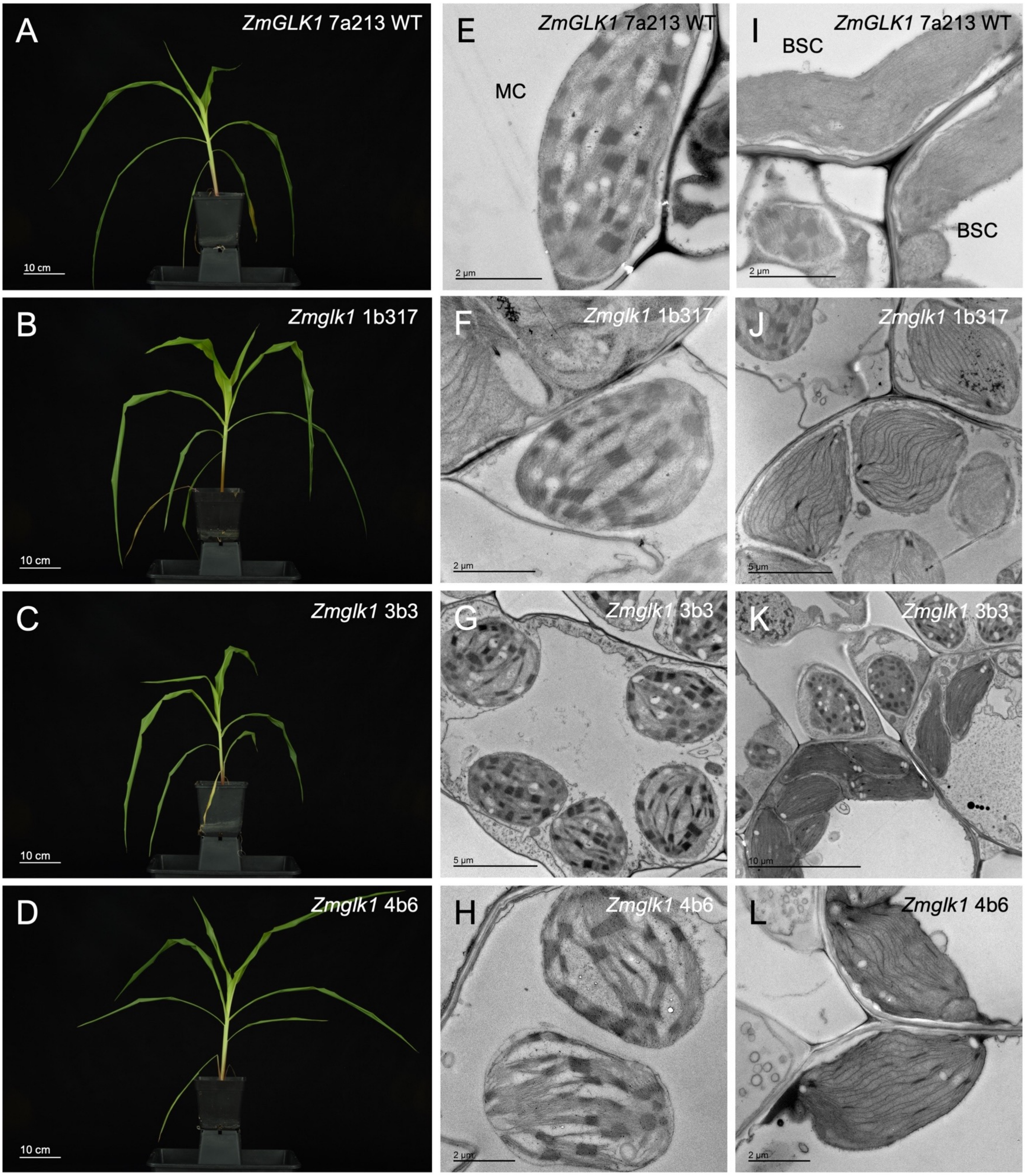


**Figure S2. Phenotypic characterization of three independent *Zmglk1* mutant lines. A-D)** Whole plant phenotype 30 days after sowing. Scale bars = 10 cm. **E-H)** Transmission electron microscopy (TEM) images showing mesophyll chloroplast ultrastructure. Scale = 2 μm, 2 μm, 5 μm and 2 μm, respectively. **I-L)** TEM images showing bundle sheath chloroplast ultrastructure. Scale bars = 2 μm, 5 μm, 10 μm and 2 μm, respectively.

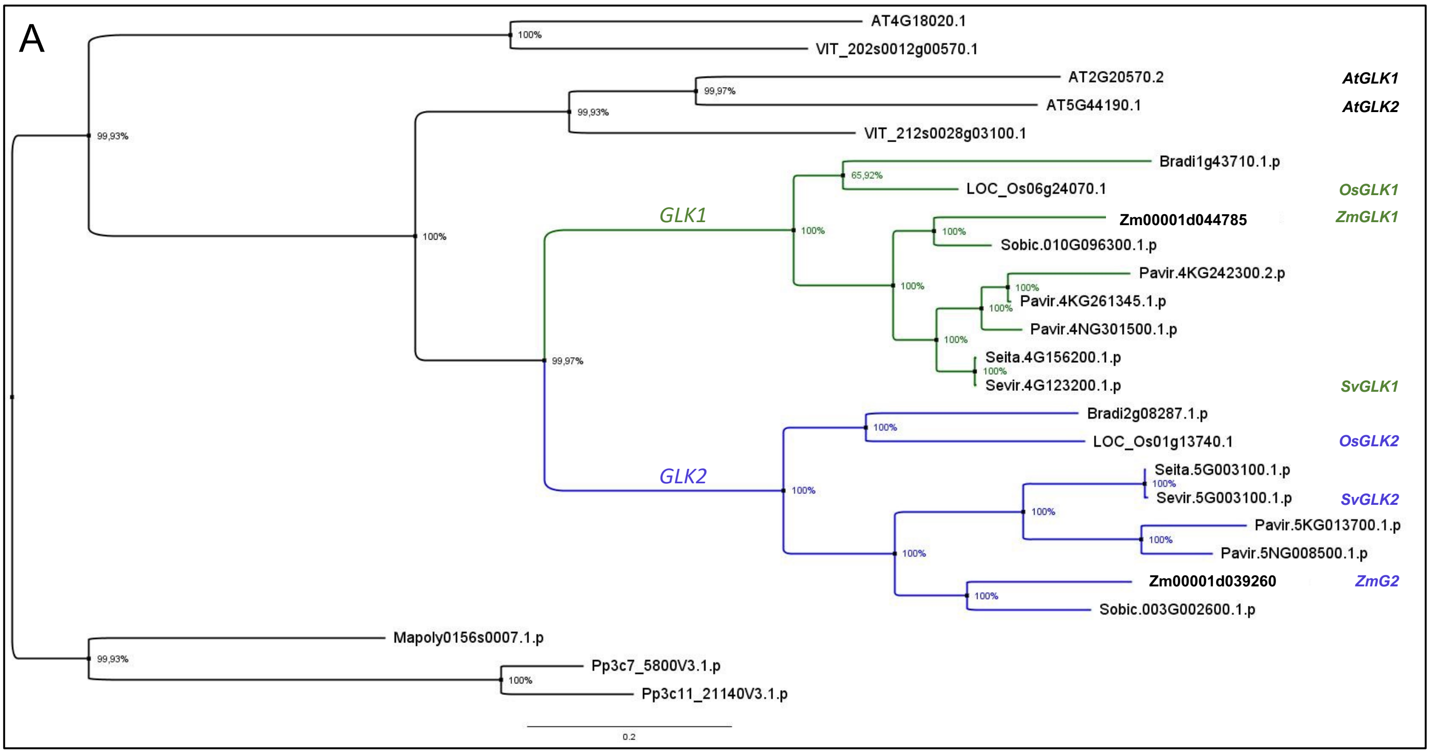


B

```
ZmGLK1      MLAVSPSPV------RCADAEECGGGGASKEMEETAVGPVSDSDLDFD---FTVDDIDFGDF--FLRLDDGDDALPGLEVDPAEIVFADFEAIATAG-----GDGGVTDQEVPSVLPFAD
SvGLK1      --MLAVSPV------RCADADECGGGAGM---EMEEIGPV--SDLDFD---FTVDDIDFGDF--FLRLEDGD-ALPDLEVDPAEI-FTDFEAIA--T-----GSDGVMDQEVPSVQPLTD
ZmG2        --MLEVSTLRGPTSSGSKAEQHCGGGGGFVGDHHVV--FPTSGDCFAMVDDNLLDYIDFS--CDVPFFDADGDILPDLEVDTTELL-AEFSSTPPADDLLAVAVFGADDQPAAAVAQE--
SvGLK2      --MLEVSSLRSPSKA--------ADQRGFVGDHVGF--PTSAGTDDFMVDDNLLDYIDFSSCCDMPFFDADGDILPDLEVDPTESLLAEFSSPE-TSPAAADGGGGAEAKPPDEKQEQVA
               :  * :                ..  .    .         .          :* ***.    .  :: ..   **.**** :*   ::*.:          .  *.  :

ZmGLK1      AAHIGAVDPCCGVLGEDNDAACADVEEGKGECDHADEVAAAGNNNSDSGEAGCGGAFAGEKSPSSTASSSQEAESRRKVSKKHSQGKKKAKVDWTPELHRRFVQAVEELGIDKAVPSRIL
SvGLK1      AVHLDALDPCSVVLGEDNTT-RADVEEGKGECNHAEEVVPGNGDF----GGGGGTVLAEEKSPSSTTSSSQEAESRHKSSSKHSHGKKKAKVDWTPELHRRFVQAVEQLGIDKAVPSRIL
ZmG2        -KPSSSLEQTC---GDDKGVAVAAARRKL--QTTTTTTT---TEEEDSSPAGSGAN---K-SS---ASAEGHSSKKKSAGKNSNGGKRKVKVDWTPELHRRFVQAVEQLGIDKAVPSRIL
SvGLK2      AKTAAAEETKA---DDDEKTAAGGVVVTR--KRNGDEVT---TTEEDSAGAGAGSD---TKSS---AAAGGHSKKKASAGKDS-NGKRKVKVDWTPELHRRFVQAVEQLGIDKAVPSRIL
                 : :  .   .:*: .  . .         ..            .* *       *    :::  .:..: . ...    **:*.*****************:************

ZmGLK1      EIMGIDSLTRHNIASHLQKYRSHRKHMLAREVEAATWTTHRRPMYAA--PSGAV--KRPDSNAWTVPTIGFPPPAG--TPPRPVQHFGRPLHVWGHPSPTPA-----VESPRVPMWPRHL
SvGLK1      EIMGIDSLTRHNIASHLQKYRSHRKHMLAREAEAASWTQRRQ-MYAAGGASTAV--KRPDSNAWTVPTIGFPPPPPPPPPPHPMQHFGRPLHVWGHPT--PG-----VESPRVPMWPRHL
ZmG2        EIMGTDCLTRHNIASHLQKYRSHRKHLMAREAEAATWAQKRH-MYAPP-APRTTTTTDAARPPWVVPTT---------I-GFPPPRFCRPLHVWGHPPPHAAAAEAAAATPMLPVWPRHL
SvGLK2      EIMGIEGLTRHNIASHLQKYRSHRKHLMAREAEAATWAQKRQ-LYAAAPVARKVEAAAAAGPPWVVPTIGFPPPPPPAM-APPPPPFCRPLHVWGHPPTAAVE----APATMLPVWPRHL
            **** : *******************::***.***:*: :*: :**        .          *.***                  *   * *********         . :  :*:*****

ZmGLK1      APRAPPPPPWAPPPPADPASFWHHAYMRQGPAA--HMPDQVAVTPCVAVPMAAARFPAPHVRGSLPWPPPMYRPLV--PPALAGKSQQDALFQLQIQPSSESIDAAIGDVLTKPWLPLPL
SvGLK1      VPRAPT-PPWAPPPPSDP-AFWHHPYMRGPA----QMPG--QMAPCMAVPMPAARFPAPPVRGVLPCPPPMYRPLV--PPVLANKSQQDAQLQLQTQPSSESIDAAIGDVLSKPWLPLPL
ZmG2        APPRHL-APWAHPTPVDP-AFWHQQYSAA-RKWGPQAAAVTQGTPCVPLPRF-----P--VPHPIYSRPAMVPPP------PSTTKLAQLHLELQAHPSKESIDAAIGDVLVKPWLPLPL
SvGLK2      APPRP----W-----VDP-GYWHQQYNAGRTKWAPPQAAVTQGTPCVPPPPAMVQLPR--FPHPMYSRPPVMVPPPPPPPTTTTTKLAELQLQLDAHPSKESIDAAIGDVLVKPWLPLPL
            .*       *      ** .:**: *                :**:  *          .   :   * :  *         : ..  :  ::*: :**.*********** ********

ZmGLK1      GLKPPSVDSVMGELQRQGVANVPQACG*-------
SvGLK1      GLKPPSVDSVMDELQRQGVADVPPACG*-------
ZmG2        GLKPPSLDSVMSELHKQGVPKIPPAAATTTGATG*
SvGLK2      GLKPPSLDSVMSELHKQGIPKIPPASSTG*-----
            ******:****.**::**: .:* *..
```

**Figure S3. Comparison of GLK sequences**. **A)** Phylogenetic reconstruction of the *GLK* gene family. Protein sequences of the extensively characterised GLK members ZmGLK1, ZmG2, OsGLK1 and OsGLK2 were used to create a HMMER profile that was used as a query to search for homologs in whole protein databases of 11 species (*Zea mays, Setaria viridis, S. italica, Panicum virgatum, Sorghum bicolor, Oryza sativa, Brachypodium distachyon, Vitis vinifera, Arabidopsis thaliana, Marchantia polymorpha* and *Physcomitrium patens*). The presence of the MYB domain in those sequences was confirmed using InterproScan, and only the sequences containing the conserved AREAEAA domain characteristic of the GLK subfamily were selected. The resulting 25 amino acid sequences were aligned using Clustal Omega. Phylogenetic reconstruction was performed using Bayesian inference on Mr. Bayes using Poisson with fixed substitution rates as a model. Two independent MCMC runs with 1,000,000 generations were performed and no further improvement of log likelihood of prior probabilities was observed. The tree was visualized on FigTree. Clade support on each node was assessed using Bayesian posterior probabilities. Members of the GLK1 subgroup are depicted in green and GLK2 in blue. **B)** Amino acid sequence alignment showing conservation between ZmGLK1, SvGLK1, ZmG2 and SvGLK2. The diagonal linesl show the HLH associated with MYB or MYB-related proteins; the AREAEAA conserved site is depicted in horizontal lines and the last exon that contains the GCT-box domain is depicted in vertical lines.

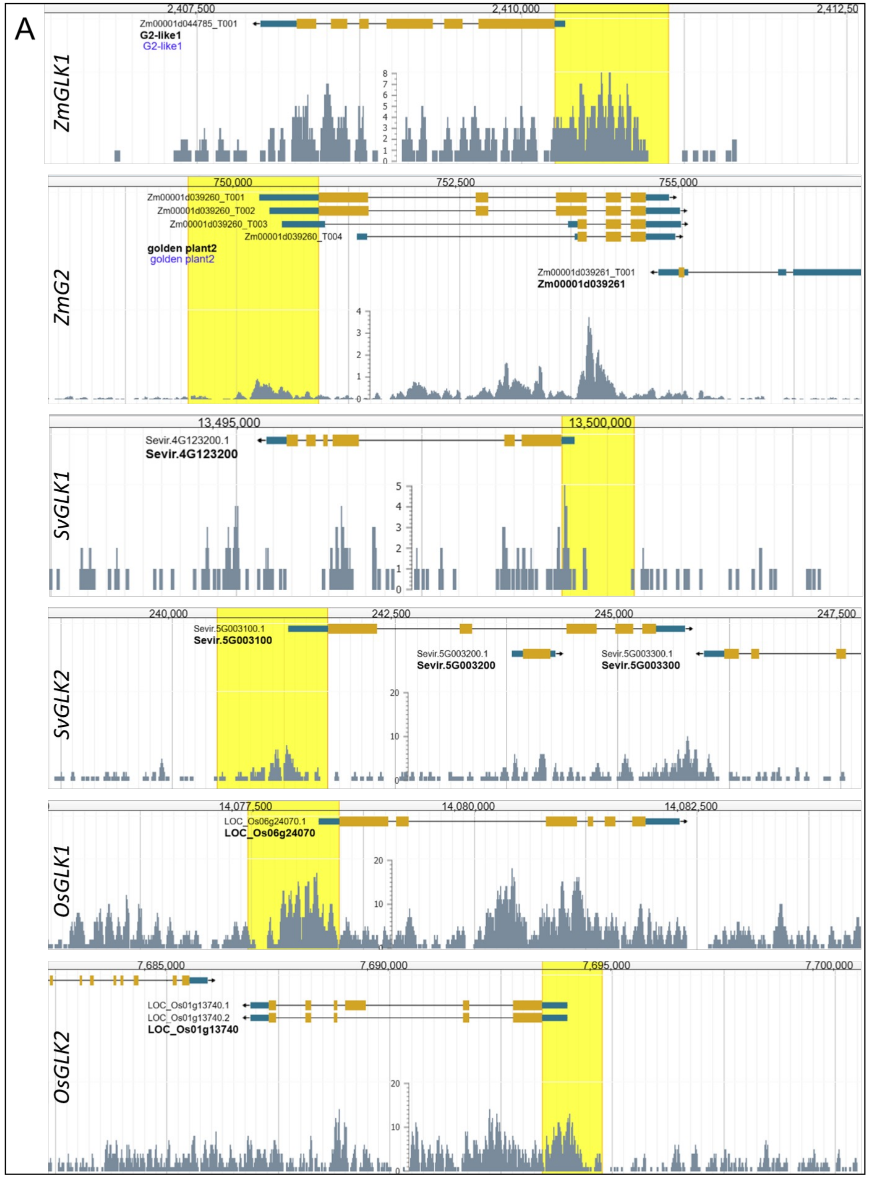


B

| Motif | ZmGLK1 | ZmG2 | SvGLK1 | SvGLK2 | OsGLK1 | OsGLK2 |
|---|---|---|---|---|---|---|
| ABI3-like | 1 | 3 | 2 | 0 | 0 | 2 |
| BPC | 1 | 47 | 1 | 81 | 19 | 133 |
| GATA | 3 | 9 | 17 | 3 | 0 | 4 |
| MYB-B | 3 | 1 | 2 | 0 | 6 | 2 |
| TGA | 1 | 0 | 0 | 0 | 0 | 0 |
| WRKY | 20 | 0 | 0 | 21 | 0 | 0 |
| bZIP/bHLH (G-box) | 0 | 8 | 1 | 3 | 0 | 8 |
| MYB-like/I-box | 0 | 10 | 10 | 3 | 9 | 7 |
| TCP | 0 | 1 | 0 | 0 | 1 | 1 |
| GT-1 | 0 | 0 | 0 | 0 | 0 | 1 |

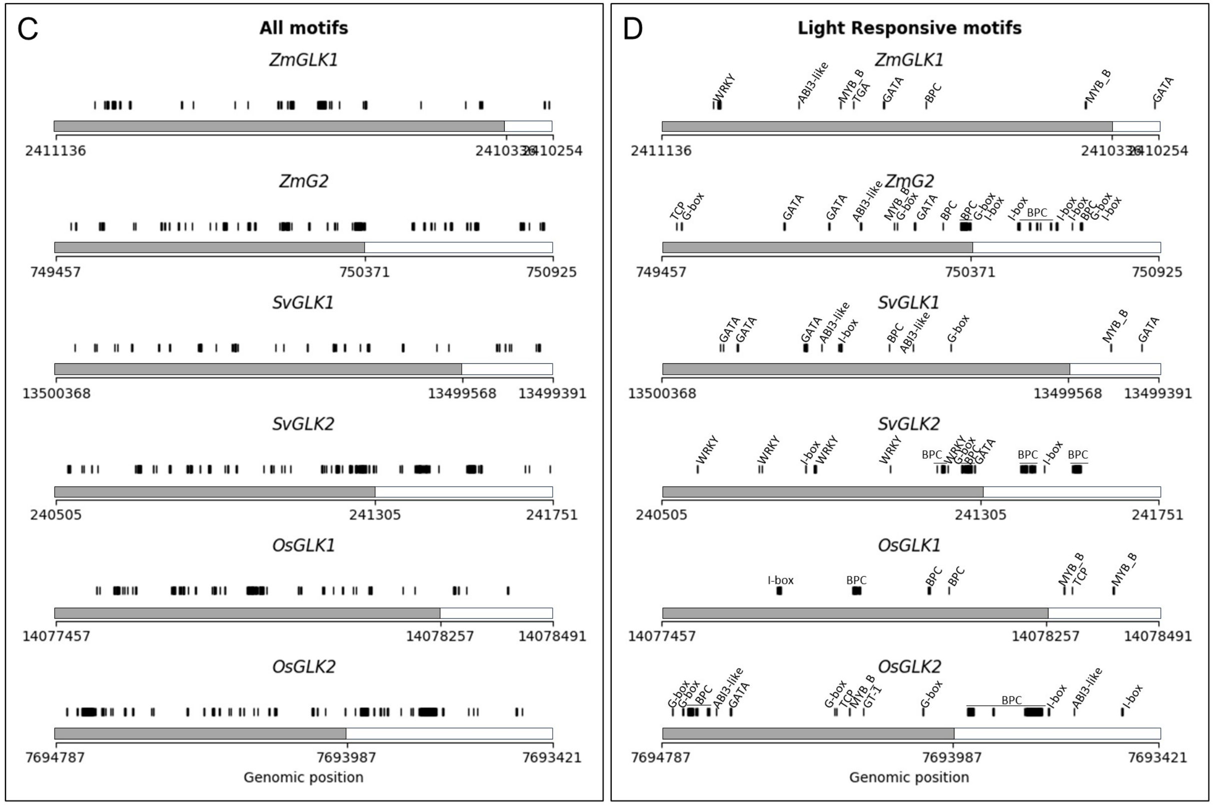


**Figure S4. DNA binding motifs in regulatory regions of *GLK* genes. A)** Schematics of maize, setaria and rice *GLK* genes showing binding sites revealed from ATAC assays with leaf RNA. Data was retrieved from https://epigenome.genetics.uga.edu/PlantEpigenome/index.html. The yellow shading highlights the 800 bp upstream of the transcription start site plus the 5'UTR regions. **B)** Table showing the number of DNA binding motifs found in regulatory regions of each gene. Plant non-redundant motifs (JASPAR) was used to screen the above-mentioned regions of each gene, using the FIMO tool with a p value of <0.0001. The photosynthesis-related/light responsive motifs (BPC, ABI3-like, GT-1, GT-4, CNA, MYB-B, STZ, bZIP/bHLH(G-box), GATA, MYB-like/I-box, TCP, TGA and WRKY) were as described by Sing et al. (2023) and rice and maize GLK-specific binding sites were as described in Tu et al. (2022). **C, D)** Schematics of 800 bp upstream of the transcription start site plus the 5'UTR regions for each gene, showing the position of all predicted DNA binding motifs (C) and of known light-responsive motifs (D). The 5' UTR is shown in white and promoter sequence in grey.

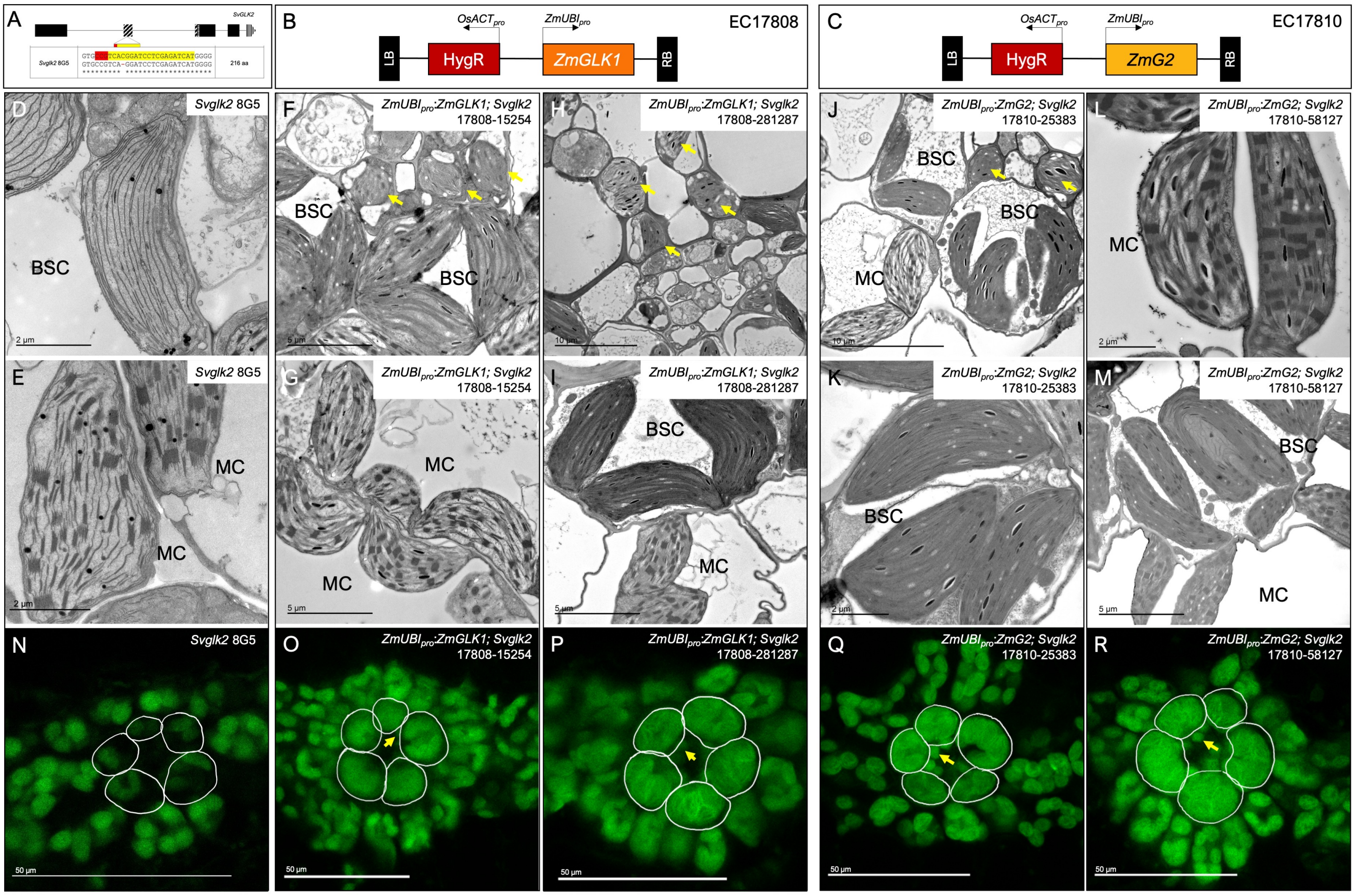


**Figure S5. Phenotypic characterization of *Svglk2* mutant lines complemented with *ZmGLK1* or *ZmG2*. A)** Schematic of the gene edited *Svglk2* mutant allele. **B-C)** Schematic of the constructs used to express *ZmGLK1* (B) or *ZmG2* (C) in the *Svglk2* background. *HygR* depicts the hygromycin phosphotransferase gene and $OsACT_{pro}$ and $ZmUBI_{pro}$ represent the constitutive rice actin and maize ubiquitin promoters, respectively. LB and RB refer to left and right borders. **D-M)** Transmission electron microscopy images showing mesophyll (MC) and bundle sheath (BSC) cell chloroplast ultrastructure in the mutant line (D, E) and in lines overexpressing *ZmGLK1* (F-I) or *ZmG2* (J-M) in the mutant background. Scale bar sizes are indicated on each image. D & E are the same images as Figure 3G & 3H). **N-R)** Confocal images of leaf cross sections from the mutant (N) and from lines overexpressing *ZmGLK1* (O, P) or *ZmG2* (Q, R) showing morphology of chloroplasts. Bundle sheath cells are outlined in white. Scale bars = 50 µm. Yellow arrows indicate ectopic chloroplast formation in vascular cells

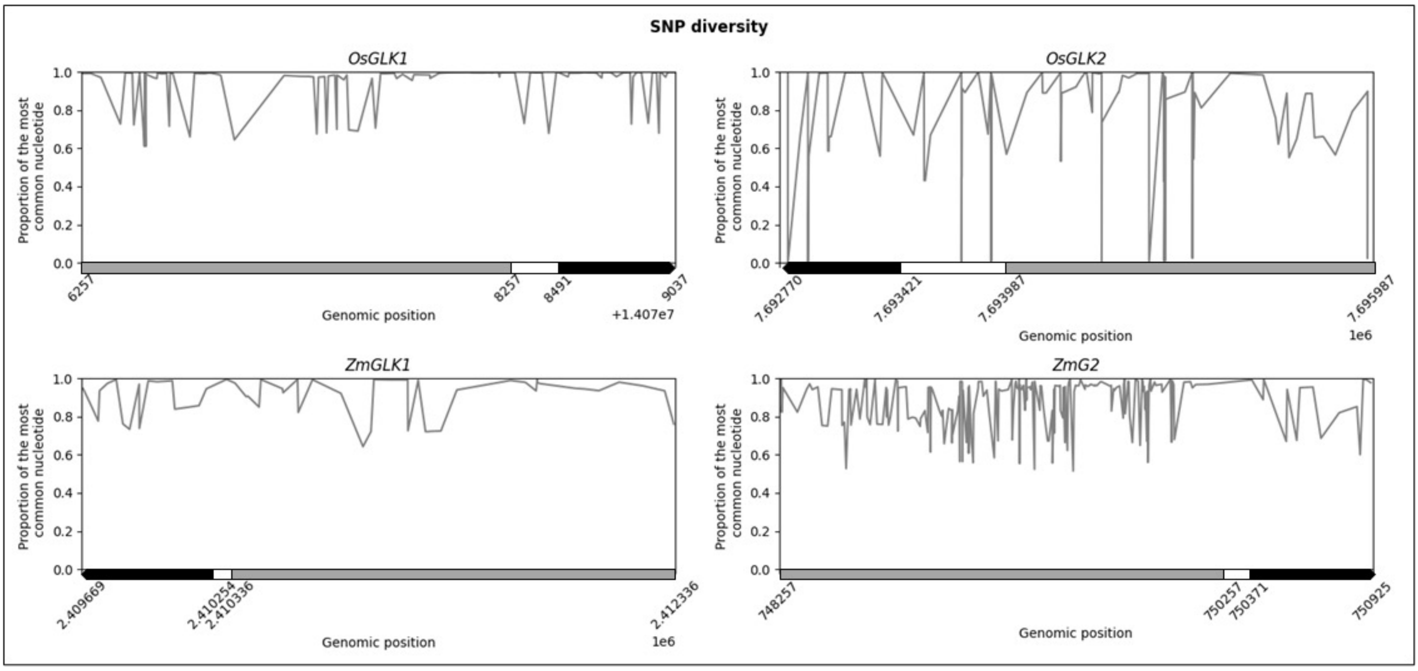


**Figure S6. Sequence variability in regulatory regions of *GLK* genes.** The SNPs from 3025 variants of rice (SNP seek, IRRI) and 1210 variants of maize (Maize SNPDB) were used to calculate the frequency of each nucleotide for each SNP position. The nucleotide with highest prevalence in each position is plotted on the graphs for *OsGLK1*, *OsGLK2, ZmGLK1* and *ZmG2*. The 2Kb upstream of the transcription start site is shown in grey, the 5'UTR in white and the first exon in black.

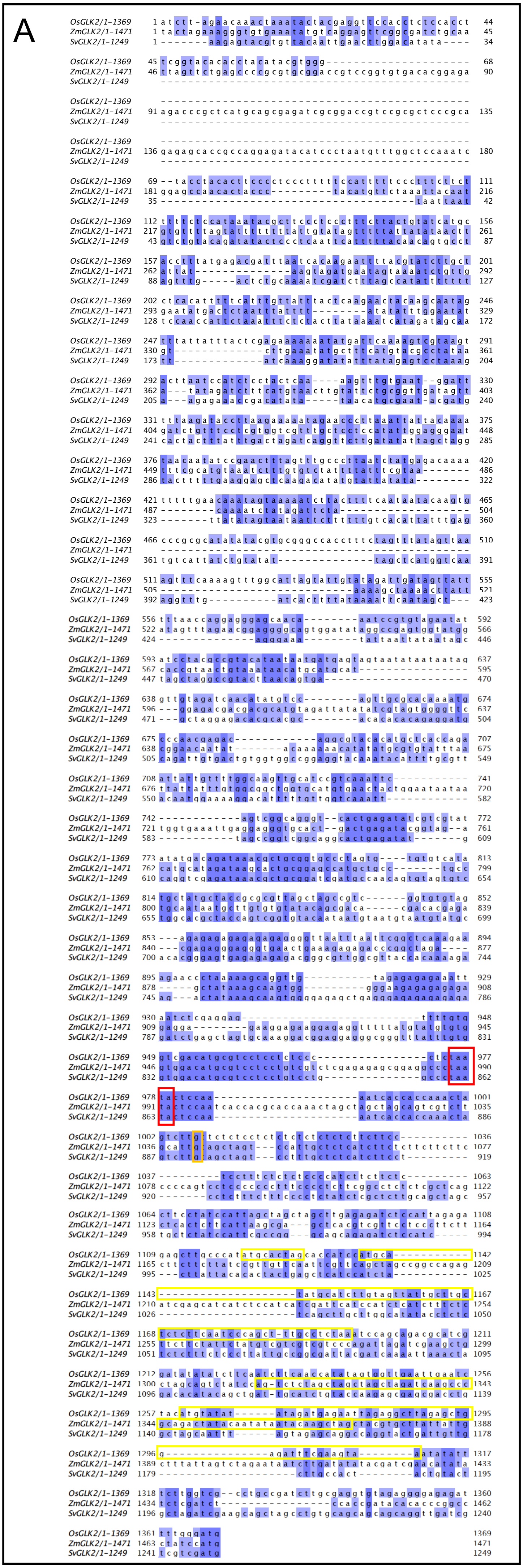


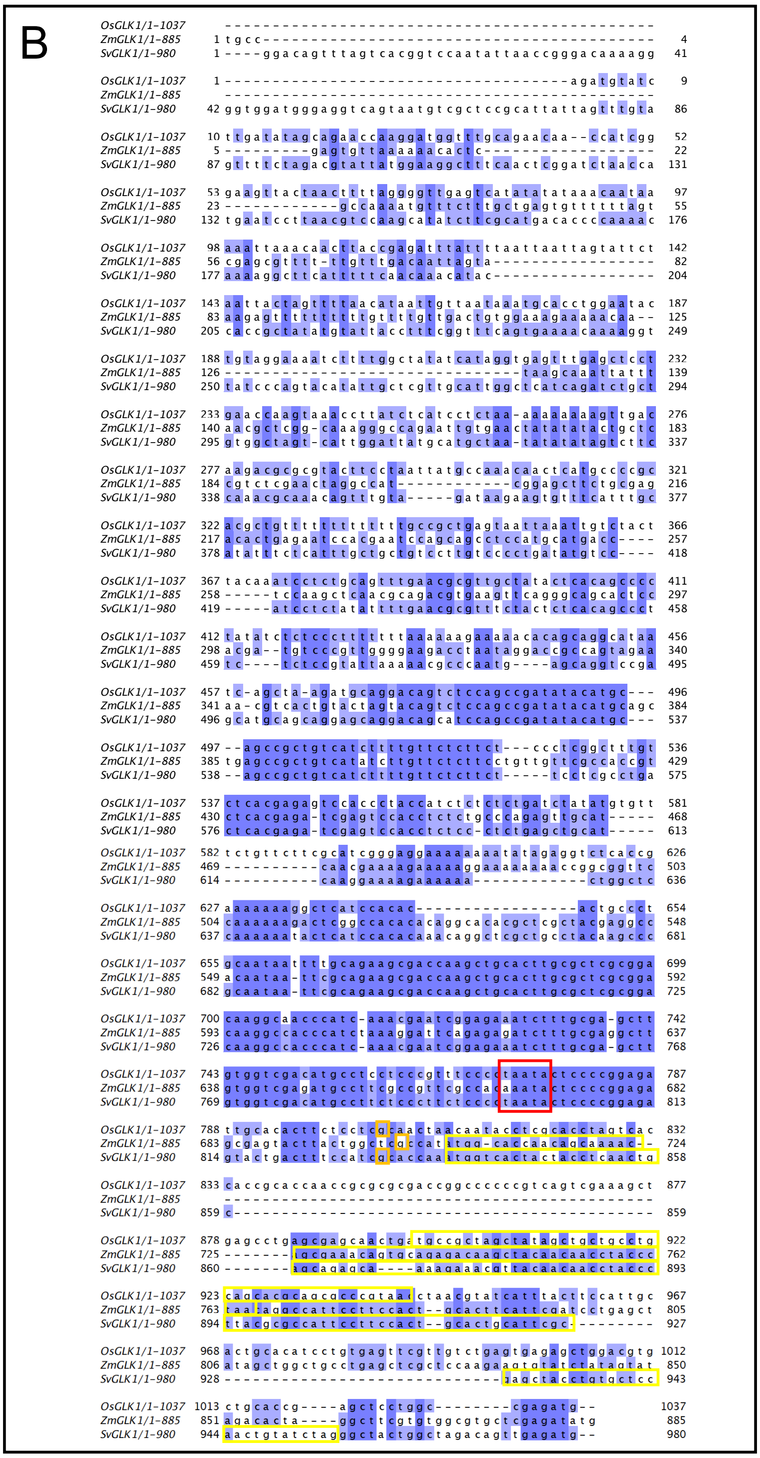


C

OsGLK1: MPLAIAAACSTQRP*

OsGLK2: MH*

MHMHLVVIACSLQSQLCL*

MRIRGLELEISK*

ZmGLK1: MAPTAKQRNSAETSYNNLP*

ZmGLK2: MSSSSQIRSKLLAVYPVSS*

SvGLK1: MVTTTSTAAEQKKRYNNLPLRAIPSTAL HSRATCAPTVSRATG*

**Figure S7. Alignment of *GLK* gene promoter and 5' UTR regions**. **A, B)** *GLK1* (B) and *GLK2* (A) sequences (800 bp upstream of the transcription start site (TSS) plus the 5' UTR) aligned using MAFFT (Katoh *et al.* 2019) and visualized using JALview (Waterhouse *et al.* 2009). Conserved bases are indicated by blue shading, the predicted TATA box by red squares, actual (rice - Murray *et al.* 2022) or predicted TSS by orange squares & putative uORFs by yellow squares. **C)** Amino acid sequences of predicted uORFs in (A) & (B).

| Primer Name | Description | F/R | Sequence(5'-3') | Amplicon gDNA | Amplicon cDNA |
|---|---|---|---|---|---|
| **Cloning oligonucleotides** | | | | | |
| JLF_Ox43 | *SvGLK1* gRNA | F | acttgTCGAAATCCAGGTCCGACACg | NA | NA |
| JLF_Ox44 | *SvGLK1* gRNA | R | aaaacGTGTCGGACCTGGATTTCGAc | | |
| JLF_Ox51 | *SvGLK2* gRNA | F | acttgATGATCTCGAGGATCCGTGAg | NA | NA |
| JLF_Ox52 | *SvGLK2* gRNA | R | aaaacTCACGGATCCTCGAGATCATc | | |
| **Genotyping primers** | | | | | |
| JLF_Ox12 | HygR | F | AGGCTCTCGATGAGCTGATGCTTT | 335 bp | NA |
| JLF_Ox13 | HygR | R | AGCTGCATCATCGAAATTGCCGTC | | |
| JLF_Ox14 | DNA control for S. viridis | F | CAGCAAGCCGCCTATATGGAG | 542 bp | NA |
| JLF_Ox15 | DNA control for S. viridis | R | TCGTCTCAGGAGTGGCCAAGT | | |
| JLF_Ox45 | CRISPR/Cas9 *SvGLK1* | F | GCTACCTGTGCTCCAACTGT | 303 bp (WT) | NA |
| JLF_Ox46 | CRISPR/Cas9 *SvGLK1* | R | CACCTCCTGGTCCATGACAC | | |
| JLF_Ox53 | CRISPR/Cas9 *SvGLK2* | F | TGTCTTGGCTCCCATATGCA | 345 bp (WT) | NA |
| JLF_Ox54 | CRISPR/Cas9 *SvGLK2* | R | TGGCTTGCAATGTTGTGTCG | | |
| JLF_Ox178 | t-DNA *ZmGLK1* | F | ATTTGGCTTGACCTTGATGG | 353 bp | NA |
| JLF_Ox179 | t-DNA *ZmGLK1* | R | GGGGATACGTACACGGTTGA | | |
| JLF_Ox175 | CRISPR/Cas9 *ZmGLK1* | R | CTTGGGAGTGCTTCTTGGAC | 458 bp (WT) | NA |
| JLF_Ox39 | CRISPR/Cas9 *ZmGLK1* | F | GGACCTGGATTTCGACTTCA | | |
| **qPCR primers** | | | | | |
| JLF_Ox26 | Si035045 (Reference gene 1) | F | AAGTTTTTGCGTCACTTCCACT | 500 bp | 139 bp |
| JLF_Ox27 | Si035045 (Reference gene 1) | R | CAAATTTCTGCCCTCGCTAAT | | |
| JLF_Ox28 | Si03461 (Reference gene 2) | F | ACGAGAAGGATTCATCCAAGAC | 195 bp | 98 bp |
| JLF_Ox29 | Si03461 (Reference gene 2) | R | CGTGTACTCTTGCATCTTGGAG | | |
| JLF_Ox103 | SvGLK2 (*Svglk1* and A10 only) | F | TCGCACAGAAAGCACCTGAT | 652 bp | 443 bp |
| JLF_Ox104 | *SvGLK2* (*Svglk1* and A10 only) | R | CTGTACATGGGGTGCGGAAA | | |
| JLF_Ox119 | *SvGLK2* | F | AGGAAGAGGAACGGGGATGA | 1086 bp | 163 bp |
| JLF_Ox120 | *SvGLK2* | R | CCAGTCCACCTTCACCTTCC | | |
| JLF_Ox151 | *SvGLK1* | F | TGTGTCGGACCTGGATTTCG | 172 bp | 172 bp |
| JLF_Ox152 | *SvGLK1* | R | CACCTCCTGGTCCATGACAC | | |
| JLF_Ox101 | *SvGLK1* (*Svglk1* and A10 only) | F | GTACCGGTCTCACAGGAAGC | 620 bp | 450 bp |
| JLF_Ox102 | *SvGLK1* (*Svglk1* and A10 only) | R | ACAAGGCAAAACACCCCTCA | | |
| ZmCYP_F | *ZmCYP* (Reference gene 1) | F | CTGAGTGGTGGTCTTAGT | 100 bp | 100 bp |
| ZmCYP_R | *ZmCYP* (Reference gene 1) | R | AACACGAATCAAGCAGAG | | |
| ZmEF1α_F | *ZmEF1α* (Reference gene 2) | F | TGGGCCTACTGGTCTTACTACTGA | 135 bp | 135 bp |
| ZmEF1α_R | *ZmEF1α* (Reference gene 2) | R | ACATACCCACGCTTCAGATCCT | | |
| JLF_Ox173 | *ZmGLK1* | F | GATCCGTGTTGTGGTGTCCT | 111 bp | 111 bp |
| JLF_Ox174 | *ZmGLK1* | R | ATTATTACCGGCGGCTGCTA | | |
| JLF_Ox37 (57RTF) | *ZmG2* | F | CATGGTGGACGACAACCTC | 236 bp | 236 bp |
| JLF_Ox38 (57RTR) | *ZmG2* | R | CACATGTTTGCTCCAACGAC | | |
| JLF_Ox197 | *ZmGNC* | F | GTGTCAGCAGTTCCCACTGA | 105 bp | 105 bp |
| JLF_Ox198 | *ZmGNC* | R | CCACGTCCACTCTCTTCTCC | | |
| JLF_Ox199 | *ZmCGA* | F | GGCAGCATGTCTTGTCTGAA | 265 bp | 119 bp |
| JLF_Ox200 | *ZmCGA* | R | CACCATCGATGCTTTGGATA | | |
